# Enhanced *in Vivo* Blood Brain Barrier Transcytosis of Macromolecular Cargo Using an Engineered pH-sensitive Mouse Transferrin Receptor Binding Nanobody

**DOI:** 10.1101/2023.04.26.538462

**Authors:** Thomas J. Esparza, Shiran Su, Caroline M. Francescutti, Elvira Rodionova, Joong Hee Kim, David L. Brody

## Abstract

**Background:** The blood brain barrier limits entry of macromolecular diagnostic and therapeutic cargos. Blood brain barrier transcytosis via receptor mediated transport systems, such as the transferrin receptor, can be used to carry macromolecular cargos with variable efficiency. Transcytosis involves trafficking through acidified intracellular vesicles, but it is not known whether pH-dependent unbinding of transport shuttles can be used to improve blood brain barrier transport efficiency.

**Methods:** A mouse transferrin receptor binding nanobody, NIH-mTfR-M1, was engineered to confer greater unbinding at pH 5.5 vs 7.4 by introducing multiple histidine mutations. The histidine mutant nanobodies were coupled to neurotensin for *in vivo* functional blood brain barrier transcytosis testing via central neurotensin-mediated hypothermia in wild-type mice. Multi-nanobody constructs including the mutant M1_R56H, P96H, Y102H_ and two copies of the P2X7 receptor-binding 13A7 nanobody were produced to test proof-of-concept macromolecular cargo transport *in vivo* using quantitatively verified capillary depleted brain lysates and *in situ* histology.

**Results:** The most effective histidine mutant, M1_R56H, P96H, Y102H_ -neurotensin, caused >8°C hypothermia after 25 nmol/kg intravenous injection. Levels of the heterotrimeric construct M1_56,96,102His_-13A7-13A7 in capillary depleted brain lysates peaked at 1 hour and were 60% retained at 8 hours. A control construct with no brain targets was only 15% retained at 8 hours. Addition of the albumin-binding Nb80 nanobody to make M1_R56H, P96H, Y102H_ -13A7-13A7-Nb80 extended blood half-life from 21 minutes to 2.6 hours. At 30-60 minutes, biotinylated M1_R56H, P96H, Y102H_ -13A7-13A7-Nb80 was visualized in capillaries using *in situ* histochemistry, whereas at 2-16 hours it was detected in diffuse hippocampal and cortical cellular structures. Levels of M1_R56H, P96H, Y102H_-13A7-13A7-Nb80 reached more than 3.5 percent injected dose/gram of brain tissue after 30 nmol/kg intravenous injection. However, higher injected concentrations did not result in higher brain levels, compatible with saturation and an apparent substrate inhibitory effect.

**Conclusion:** The pH-sensitive mouse transferrin receptor binding nanobody M1_R56H, P96H, Y102H_ may be a useful tool for rapid and efficient modular transport of diagnostic and therapeutic macromolecular cargos across the blood brain barrier in mouse models. Additional development will be required to determine whether this nanobody-based shuttle system will be useful for imaging and fast-acting therapeutic applications.

## BACKGROUND

The delivery of biological macromolecular cargos across the blood brain barrier (BBB) is a major unmet need for diagnostic and therapeutic approaches to brain disorders. In recent years, it has become clear that taking advantage of endogenous receptor mediated transcytosis systems represents a promising approach. To that end, several research groups have developed antibodies and other reagents that bind key targets involved in receptor mediated transcytosis in brain endothelial cells [1] [2]. A common target is the transferrin receptor (TfR) [3] [4], involved in the transcytosis of iron carried by transferrin across the BBB, where it is required for multiple iron-containing proteins in multiple brain cell types [5]. The first clinically approved therapeutic employing a receptor mediated transcytosis BBB shuttle, pabinafusp alfa, consists of a TfR-binding monoclonal antibody fused to the enzyme iduronate-2-sulfatase, which is defective in mucopolysaccharidosis type II (Hunter syndrome) [6] [7]. Additional targets for BBB transcytosis shuttles include the insulin receptor [8] [9], IGF1 receptor [10] [11] [12], CD98hc [13] [14], TMEM30 [15], and others [1] [2] [16].

To date, the majority of the candidate BBB transcytosis shuttles have been designed for long-term therapeutic applications, but there remains an unmet need for systems to shuttle biological macromolecular cargos across the BBB optimized for rapid molecular contrast imaging and fast-acting therapeutics. Furthermore, shuttle systems that are readily engineered, inexpensive to produce, simple to handle, and widely accessible to the research community would be a benefit. For specific applications to fast-acting therapeutics and imaging, rapid BBB kinetics would be an advantage. Nanobodies, 12-15 kDa recombinant proteins derived from the binding domains of heavy chain only antibodies from camelids [17] [18], have potential to address this unmet need. Nanobodies are relatively small, readily engineered, and inexpensive to produce. Because of their small size, nanobody kinetics may be expected to be relatively fast [19], though they can be engineered to have prolonged half-lives in blood [18]. The first approved nanobody therapeutic, caplacizumab-yhdp which targets von Willebrand factor, has proven to be effective for the rapid treatment of thrombotic thrombocytopenic purpura [20] [21] [22] [23]. Additional nanobody-based therapeutics approved for human use include ciltacabtagene autoleucel which targets B-cell maturation antigen for multiple myeloma [24], envafolimab which targets programmed death ligand 1 for solid tumors [25] and ozoralizumab which targets tumor necrosis factor alpha for rheumatoid arthritis [26]. Other researchers have developed nanobody-based BBB transcytosis reagents that bind to TMEM30 [15], TfR [27], [28], and IGF1R [10] [11].

We previously reported the generation and characterization of a llama nanobody recognizing the extracellular domain of the mouse transferrin receptor called NIH-TfR-M1, referred to as M1 herein. We showed that this nanobody binds independently of the presence or absence of mouse holo-(iron bound) transferrin with relatively high (5 nM) affinity [29]. The M1 nanobody showed modest BBB transcytosis *in vivo* as measured using a neurotensin (NT)-induced hypothermia assay [29]. Others have reported that antibodies with lower affinities or monovalent binding mediate improved BBB transcytosis [30] [31] [32]. It has also been reported that at least some of the intracellular vesicles involved in receptor mediated transcytosis are acidified to approximately pH 5.5 [5], and that pH-dependent binding may favor TfR antibody transcytosis *in vitro* [33]. Of note, iron dissociates from transferrin under acidic conditions due to protonation of transferrin histidine residues, and apo-transferrin has a much lower affinity for TfR than holo-transferrin [5]. Thus, we reasoned that introduction of histidine residues into M1 could improve BBB transcytosis via accelerated unbinding from TfR at acidic pH while retaining relatively fast binding kinetics at pH 7.4. We were inspired by a previous approach taken using histidine engineering of a TfR-binding single chain variable fragment (scFv) to enhance intracellular accumulation [34], as well as previously reported pH-dependent binding of candidate shuttles to other BBB transport systems [10] [14]. Researchers have introduced histidine mutations for pH-dependent engineering in many other antibodies for other purposes [35] [36] [37]. As an initial foray, we previously reported that introduction of a single histidine mutant at residue 96 imparts enhanced unbinding at pH 5.5 and modestly improved *in vivo* BBB transcytosis as measured using the NT-induced hypothermia assay. However, the M1_P96H_ mutant nanobody was still not effective as a carrier of amyloid-beta binding nanobodies across the BBB in a mouse model of Alzheimer Disease-related amyloid plaque pathology [29]. Therefore, we endeavored to further engineer the M1 nanobody for more extensive pH-dependent unbinding while retaining high affinity at pH 7.4. To this end, we produced 39 additional histidine mutant versions of M1, and characterized them both *in vitro* and *in vivo*. One version with 3 histidine mutations, M1_R56H, P96H, Y102H_, demonstrated nearly 3-fold greater unbinding at pH 5.5 compared with pH 7.4 *in vitro*, greater than 24-fold stronger NT-induced hypothermia effects than M1_P96H_, and capacity to transport more than 3.5% of the injected dose of a proof-of-concept biological macromolecule cargo per gram of brain across the BBB *in vivo* at 4 hours after intravenous injection. This triple histidine mutant TfR-binding nanobody may have potential as a BBB transcytosis shuttle for fast-acting therapeutics and molecular contrast imaging applications.

## METHODS

### Development of mouse transferrin receptor binding nanobodies

The production and development of the lead candidate nanobody NIH-TfR-M1 was described previously [29]. Briefly, an adult llama was immunized with a recombinant protein consisting of the extracellular domain of the mouse transferrin receptor. Post-immune B cell lymphocyte DNA was used to generate a phage display library. Panning the phage display library using immobilized recombinant mouse transferrin receptor extracellular domain yielded several candidate nanobodies including M1, 4E6 and 1E5. Characterization of 4E6 and 1E5 will be reported separately.

### Expression in E. coli and purification

Expression of 6xHis tagged nanobodies in *E. coli* was described previously [29]. Briefly, nanobody constructs under the control of the lacZ promotor and an N-terminal pelB periplasmic translocation sequence were cloned into the pHEN2 phagemid vector and transformed into TG-1 competent E. coli. Sequence-confirmed clones were grown to mid-logarithmic phase (optical density = 0.6) and expression was induced by addition of 1mM isopropyl-β-d thiogalactoside (IPTG) followed by overnight expression at 30°C and 250rpm. The periplasmic compartment was gently released by osmotic shock and the nanobody construct purified using HisPur™ Ni-NTA resin. Endotoxin was removed using High-Capacity Endotoxin Removal Resin (Pierce) and confirmed to be <0.5 endotoxin units/mg of total protein. The constructs were polished with size exclusion chromatography (SEC) using a Superdex™75 GL 10/300 column, injected in a maximum volume of 1-mL and eluted at 1mL/min at room temperature into a 96-well collection block for fractionation. Typical final yield was 1 to 10 mg/100 mL culture of purified nanobody.

### Transferrin receptor ELISA with pH-dependent wash steps

Plate-based measurements of nanobody construct affinities were performed as described previously [29]. Briefly, 96-well plates were coated with 2 µg/mL recombinant mouse transferrin receptor extracellular domain (amino acids 89-763) overnight at 4°C, followed by blocking with 1% (w/v) bovine serum albumin (BSA) in 1x Phosphate-Buffered Saline (PBS) for 1 hour and incubated for 1 hour at room temperature with varying concentration of nanobody constructs. To assess pH-dependent unbinding, three washes were performed at either pH 7.4 or pH 5.5 for 10 minutes each. Then bound nanobody constructs were detected with peroxidase conjugated goat anti-alpaca VHH domain specific antibody (#128-035-232, Jackson Immuno Research) followed by colorimetric development with tetramethylbenzidine measured at 650 nanometer absorbance on a Biotek plate reader.

### Site-directed mutagenesis

Site-directed mutagenesis was performed as described previously [29]. Briefly, single histidine point mutations were generated using the Q5 Site-Directed Mutagenesis kit (New England Biolabs) with DNA oligonucleotides incorporating the mutation. The introduction of the point mutation was confirmed by Sanger sequencing before proceeding to protein expression and purification. For instances of multiple histidine point mutations, the entire nanobody fragment was synthesized (Twist Bioscience) and cloned into the pHEN2 phagemid for expression and purification as described above.

### Mice

All animal studies were performed in accordance with guidelines established by the National Institutes of Health for the care and use of laboratory animals (NIH publication 85-23, revised 2011) and approved by the National Institute of Neurological Disorders and Stroke (NINDS)/ National Institute on Deafness and Other Communication Disorders (NIDCD) Animal Care and Use Committee in the National Institutes of Health (NIH) Clinical Center (Protocol Number: 1406–21). Adult male and female C57BL/6J mice were obtained from Jackson labs (stock# 000664) at 8 weeks of age. They were housed in at 4-5 mice per cage for at least 1 week before starting experiments. For terminal experiments, mice were deeply anesthetized with 5% isoflurane mixed with medical air and euthanized by transcardiac perfusion with ice cold heparinized saline.

### NT construct fabrication

Nanobody constructs including the NT peptide were constructed as previously described [29]. Briefly, DNA constructs including nanobodies, (Gly-Gly-Gly-Ser)_3_ linkers, and the 13 amino acids encoding the NT peptide (ELYENKPRRPYIL) were cloned in-frame, expressed by IPTG induction, and purified as described above. Typical final yield was 1 to 10 mg/100 mL culture of purified nanobody.

### *In vivo* assessment of NT-induced hypothermia

Assessments of NT-induced hypothermia were performed as described [29]. Briefly, adult male and female C57BL/6J mice were weighed, abdominal fur was removed, and tails were warmed with a heating pad while mice were under isoflurane anesthesia. Investigators blinded to the identity of the injected constructs randomly assigned mice to injection with one of several constructs or controls between 0600 and 1200. Each construct was tested in at least 3 mice. Temperature measurements were obtained using an infrared thermometer applied to the abdomen while the mice were placed briefly under isoflurane anesthesia. Baseline temperatures were recorded serially prior to injection to ensure stability. Tail veins were injected with approximately 10 microliters per gram body weight of nanobody constructs using a 30 gauge needle in PBS at pH 7.4. Temperature measurements were obtained every 20-30 minutes for up to 180 minutes. Mice were allowed to recover from anesthesia between temperature measurements and observed for adverse effects. Individual mice were used for up to 3 experiments separated by at least 7 days. We have observed that after 3 experiments, hypothermia responses became inconsistent, even when using previously well-characterized NT constructs. Therefore, individual mice were not used for more than 3 experiments.

### Multi-nanobody construct fabrication and characterization

Construct fabrication was performed as described previously [29]. DNA sequences for the 13A7 nanobody and the Nb80 nanobody were generated based on *E. coli* preferred codon usage. Nanobody constructs included (Gly-Gly-Gly-Ser)_3_ linkers. Size and purity were assessed using size exclusion chromatography. No aggregation or degradation products were observed.

### Nanobody biotinylation

The incorporation of biotin moieties for downstream detection of nanobody constructs was achieved using a 5-fold molar excess of EZ-link-NHS-PEG4-biotin (#21330, Thermo Fisher) with nanobody constructs in 50mM sodium carbonate buffer, pH 8.5 for 1 hour at room temperature. The unreacted biotin reagent was removed using a 5-mL desalting column on an AKTA Pure FPLC system.

### Blood and brain tissue sampling

Following tail-vein injection of nanobody constructs, mice were placed under 5% isoflurane anesthesia at the specified terminal time points. Under maintained anesthesia, the right atrium was cut and approximately 100 microliters of whole blood collected into heparinized tubes. Subsequently, a 23-gauge needle was used to perfuse with heparinized ice-cold normal saline through the left ventricle. Following complete perfusion, the whole brain was extracted and bisected into separate hemispheres. The left hemisphere was placed into 4% (w/v) paraformaldehyde overnight at 4°C. The right hemisphere was snap frozen on dry ice prior to storage at -80°C for downstream capillary depleted homogenate preparation. Following 24-hour fixation of the left brain hemisphere, fixative was replaced with 30% sucrose and allowed to equilibrate at 4°C for 48-hours prior to sectioning the tissue using a freezing sliding microtome producing 50µm thick sections stored in 1xPBS containing 0.05% sodium azide prior to histological assessment.

### Capillary depletion of brain lysates

The parenchymal fraction was physically separated from the associated vessel components using a modification of previously published methods [38]. Briefly, brain hemispheres were weighed and then dounce homogenized in 1 mL ice-cold HEPES-buffered saline (HBSS) with a fixed number of dounces within sample set to reduce variability. The homogenate was transferred to a 1.5 mL microcentrifuge tube and spun at 2,000x g for 10 minutes at 4°C. The parenchymal fraction supernatant was transferred to a new tube, snap frozen on dry ice and stored at -80°C until assessed for nanobody concentration by ELISA. The vessel containing pellet was resuspended in 1mL of HBSS containing 18% (w/v) dextran (molecular weight 60-70 kDa) using a wide-bore pipet, and then spun at 4,400x g for 15 minutes at 4°C. The myelin layer was carefully removed using a disposable cotton applicator and a wide-bore pipet. Then the underlying vessel-enriched pellet was resuspended in HBSS containing 1% (w/v) BSA. The solution was transferred to a 40µm cell strainer, and the vessels were washed twice with HBSS containing 1% (w/v) BSA, followed by transfer of the vessels into a new microcentrifuge tube using a wide-bore pipet. The vessels were spun at 2,000x g for 5 minutes at 4°C, and the supernatant was removed to deplete the BSA content. The pellet was then resuspended in 1mL HBSS and spun down again at 2,000x g for 5 minutes at 4°C. The supernatant was carefully removed, and the pellet was then snap frozen on dry ice and stored at -80°C.

### *In situ* histochemistry

The presence of biotinylated nanobody construct *in situ* was performed using streptavidin-peroxidase detection followed by an amplification step as follows. Gentle antigen retrieval was performed by incubation at 60°C in 50mM sodium citrate buffer, pH 6.0 for 10 minutes. Endogenous peroxidase was then quenched by incubation in 3% (v/v) hydrogen peroxide for 10 minutes. The tissue was then blocked with 5% (v/w) BSA in 1xPBS containing 0.05% Tween-80 for 1 hour at room temperature. An initial avidin-peroxidase step was performed using Vectastain Elite ABC-HRP (#PK-6100, Vector Laboratories) reagent diluted 1:400 in 1xPBS (complexed 20 minutes before use) for 1 hour at room temperature. The biotin signal was then amplified by addition of a 2µM solution of tyramide-biotin reagent (#92176, Biotium) in 10mM Tris, pH 7.5 containing 0.0015% (v/v) hydrogen peroxide for 10 minutes at room temperature. A final avidin-peroxidase step was performed for 1 hour at room temperature. Chromogenic development was performed using diaminobenzidine. The same development termination time was used for all tissues in a sample set to provide for comparative assessments. Tissue was washed three times for five minutes each in 1xPBS between each of the above steps.

### *Ex vivo* Immunohistochemisty

Detection of *ex vivo* nanobody construct immunoreactivity in 50µm thick coronal brain sections was performed as follows: Gentle antigen retrieval was performed by incubation at 60°C in 50mM sodium citrate buffer, pH 6.0 for 10 minutes. Endogenous peroxidase was then quenched by incubation in 3% (v/v) hydrogen peroxide for 10 minutes. The tissue was then blocked with 5% (v/w) BSA in 1xPBS containing 0.05% Tween-80 for 1-hour at room temperature. Biotinylated nanobody construct was diluted to 50 nM in 0.5% (w/v) BSA in 1xPBS and tissue incubated overnight at 4°C. Avidin-peroxidase detection was performed using Vectastain Elite ABC-HRP, prepared as described above, for 1 hour at room temperature. Chromogenic development was performed using diaminobenzidine, the same development termination time was used for all tissues in a sample set to provide for comparative assessments. Tissue was washed three times for five minutes each in 1xPBS between each of the above steps.

### Percent injected dose per gram of brain calculation

Calculations were performed using the following formula:

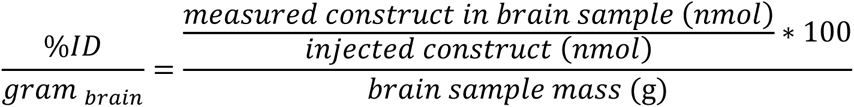

where

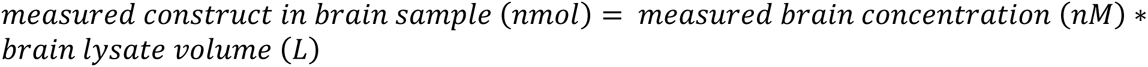

and

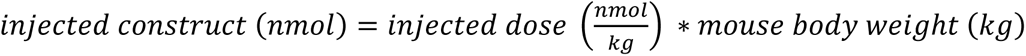

### Statistical Analyses

Statistical analyses were performed using GraphPad Prism 9.5.0. Affinity measurements were calculated using the “Dose Response” function in the “Nonlinear Regression” analysis group. Simple two way ANOVAs rather than repeated measures ANOVAs were performed because data at each time point was taken from different groups of animals. Statistical significance was defined as 2-sided p-values <0.05 after correction for multiple comparisons. Sidak’s multiple comparisons test was performed for post-hoc pairwise comparisons.

### Model fitting

GraphPad Prism 9.5.0 was used for model fitting. The models used were “Specific binding with Hill slope” and “Substrate inhibition” in the “Nonlinear Regression” analysis group. Both models had 3 free parameters and were fit to 22 pairs of injected dose vs. brain lysate concentration data points, leaving 19 degrees of freedom. Fit parameters were obtained using least squares.

## RESULTS

### Engineering the M1 mouse TfR binding nanobody with histidine mutations for pH-dependent binding

Our goal was to engineer a mTfR-binding nanobody for use as a BBB transcytosis shuttle. We refined our previously reported lead candidate llama nanobody recognizing the extracellular domain of the mouse TfR called M1 by introducing a series of histidine mutations and assessing their effects on pH dependent binding *in vitro*. We used a plate-based assay to assess relative unbinding at pH 5.5 vs. pH 7.4. In this assay, several concentrations of each M1 variant were bound to 96 well plates coated with mTfR at pH 7.4 and room temperature. Relative apparent mTfR unbinding was assessed by measuring the nanobody remaining after three 10 minute washes at either pH 7.4 or pH 5.5 (**Figure 1**). Unbinding of the original M1 was similar after three 10 minute room temperature washes at pH 7.4 vs. pH 5.5, indicating a lack of pH dependence. In contrast, several of the M1 histidine mutant had >2-fold greater unbinding at pH 5.5 vs pH 7.4. We did not observe any histidine mutants with reduced unbinding at pH 5.5. Several histidine mutants imparted pH-dependent unbinding but had reduced overall affinity. Other histidine mutants did not impart pH-dependent unbinding. The mutants, their affinities at pH 7.4, and their relative unbinding at pH 5.5 vs pH 7.4 are shown in **Table 1**. The effects of multiple histidine mutants were not necessarily additive; double, triple, quadruple, and quintuple mutants did not necessarily have greater pH-dependent unbinding than otherwise similar mutants with smaller numbers of histidines. One mutant retained high affinity and imparted steeply pH-dependent unbinding. Specifically, M1_R56H, P96H, Y102H_ had 2.63-fold greater unbinding at pH 5.5 vs pH 7.4 while retaining affinity <10 nM at pH 7.4. Exemplar binding curves for the original M1, the previously reported M1_P96H_ and the triple histidine mutant M1_R56H, P96H, Y102H_ are shown in **Figure 1**. Thus, histidine mutations introduced a range of enhanced unbinding rates at pH 5.5 *in vitro,* and one triple histidine mutant imparted both enhanced pH-dependent unbinding with preserved high affinity at pH 7.4.

**Figure 1:**
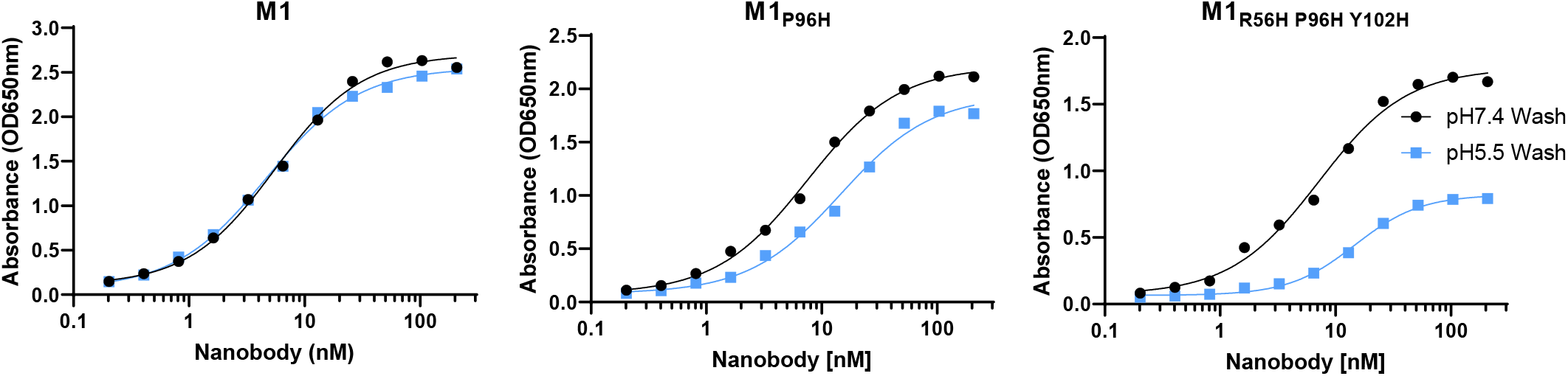
Binding and pH-dependent unbinding of M1 nanobody and histidine mutants to recombinant mTfR apical domain. Each nanobody was bound at multiple concentrations to 96 well ELISA plates coated with recombinant mTfR at pH 7.4, then washed at either pH 7.4 or pH 5.5. The original M1 has very little pH dependent unbinding, whereas the M1_P96H_ mutant had an average of 1.46-fold more unbinding at pH 5.5 and the M1_R56H, P96H, Y102H_ mutant had an average of 2.63-fold more unbinding at pH 5.5.

**Table 1:** Mouse TfR binding nanobody histidine mutants tested *in vitro* and *in vivo*. *In vitro* assessments included a) ELISA-based affinity measurements vs. recombinant mTfR extracellular domain conducted at pH 7.4 at room temperature, and b) plate-based unbinding assays conducted by binding nanobodies to recombinant mTfR extracellular domain at pH 7.4, then washing at either pH 5.5 or pH 7.4 at room temperature. *In vivo* assessments consisted of blinded measurements of mouse peak hypothermia following iv injection of 25 nmol/kg of each of the nanobodies fused with neurotensin (NT) (n=3-6 mice per construct). ND: not determined due to low binding.

| Nanobody | mTfR affinity<br>pH 7.4 (nM) | Apparent mTfR unbinding<br>ratio at pH 5.5 vs pH 7.4 | NT-mediated<br>Hypothermia (°C) |
| --- | --- | --- | --- |
| M1 | 5.11 | 1.01 | 0 |
| M1: P96H | 7.031 | 1.46 | 1 |
| M1: Y98H | 2509 | 1.28 | 2.5 |
| M1: S99H | >5000 | 1.01 | 1 |
| M1: T100aH | 55.31 | 1.05 | 1.5 |
| M1: T100bH | 30.12 | 1.24 | 0 |
| M1: R100dH | 1294 | 1.01 | 0 |
| M1: L100eH | 203 | 1.32 | 0 |
| M1: D100fH | >5000 | 1.58 | 0 |
| M1: S100gH | >5000 | 1.03 | 0 |
| M1: K100iH | >5000 | 1.42 | 0 |
| M1: D101H | 25 | 1.32 | 1.5 |
| M1: Y32H, P96H | 373 | 2.57 | 0 |
| M1: G65H, P96H | >5000 | 2.09 | 1 |
| M1: P96H, Y98H | 33.18 | 2.48 | 1.2 |
| M1: P96H, S100H | 1140 | 1.02 | 0 |
| M1: P96H, D101H | 242 | 2.65 | 7.5 |
| M1: P96H, Y102H | 11.65 | 2.33 | 6 |
| M1: Y98H, Y102H | >5000 | 1.29 | 5 |
| M1: D31H P96H Y102H | 95 | 1.13 | 0 |
| M1: R33H P96H Y102H | 1545 | 1.10 | 0 |
| M1: R56H, P96H, Y98H | >5000 | 0.98 | 1.5 |
| M1: R56H, P96H, D101H | >5000 | 1.78 | 0 |
| M1: R56H, P96H, Y102H | 6.653 | 2.63 | 8 |
| M1: R56H, Y98H, D101H | 52.6 | 1.04 | 0 |
| M1: R56H, Y98H, Y102H | >5000 | 0.99 | 0 |
| M1: R56A, P96H, Y102H | 1.7 | 2.59 | 0 |
| M1: Y59H, P96H, Y102H | 888 | 2.07 | 1 |
| M1: A60H, P96H, Y102H | >5000 | ND | 0 |
| M1: P61H, P96H, Y102H | 148 | 2.13 | 1 |
| M1: S62H, P96H, Y102H | >5000 | 2.18 | 1.5 |
| M1: A64H, P96H, Y102H | >5000 | 1.79 | 0 |
| M1: S70H, P96H, Y102H | 78 | 1.64 | 0 |
| M1: P96H S99H D101H | 301 | 2.43 | 2 |
| M1: P96H S99H Y102H | >5000 | 2.52 | 2.5 |
| M1: P96H, D101H, Y102H | >5000 | 2.42 | 0 |
| M1: C22A C92A P96H D101H | 35.76 | 1.03 | 0 |
| M1: C22A C92A P96H Y102H | 17.56 | 1.07 | 0 |
| M1: R56H, P96H, Y98H, Y102H | >5000 | 1.00 | 0 |
| M1: R56H, P96H, S99H, Y102H | 246 | 2.04 | 5.5 |
| M1: Y32H, R56H, P96H, S99H, Y102H | >5000 | 1.11 | 0 |

### *In vivo* BBB transcytosis assessed via neurotensin-induced hypothermia

We next asked whether these histidine mutants conferred greater BBB transcytosis *in vivo* using NT-induced hypothermia assays. This assay has previously been used to efficiently screen candidate BBB crossing shuttle systems [10] [27] [28] [29] [39] [40] [41]. After intravenous injection of as little as 25 nmol/kg of M1_R56H, P96H, Y102H_ -NT fusion, mouse temperatures dropped profoundly by up to 8°C (**Figure 2**). Other histidine mutants with a range of affinities also had substantially more potent effects than the original M1 or M1_P96H_ with temperature drops of 2-6 °C after intravenous injection of 25 nmol/kg (**Table 1**). In contrast, M1 and M1_P96H_ required higher doses to produce detectable drops in body temperature (**Figure 2**). An M1 variant with alanine mutations at residues 100b and 100c (M1_AA_) which did not bind TfR also did not cause detectible hypothermia at doses as high as 600 nmol/kg (**Figure 2**). M1 mutants with very low affinity generally did not produce substantial hypothermia, nor did mutants with relatively modest pH-dependent binding (**Table 1, Suppl. Fig. 1**). As noted, the NT-induced hypothermia assays were performed in a randomized, blinded fashion. These results indicated that M1 histidine mutants with pH-dependent unbinding could produce potent NT-induced hypothermia, consistent with the hypothesis that they shuttle NT across the BBB more efficiently *in vivo*.

**Figure 2:**
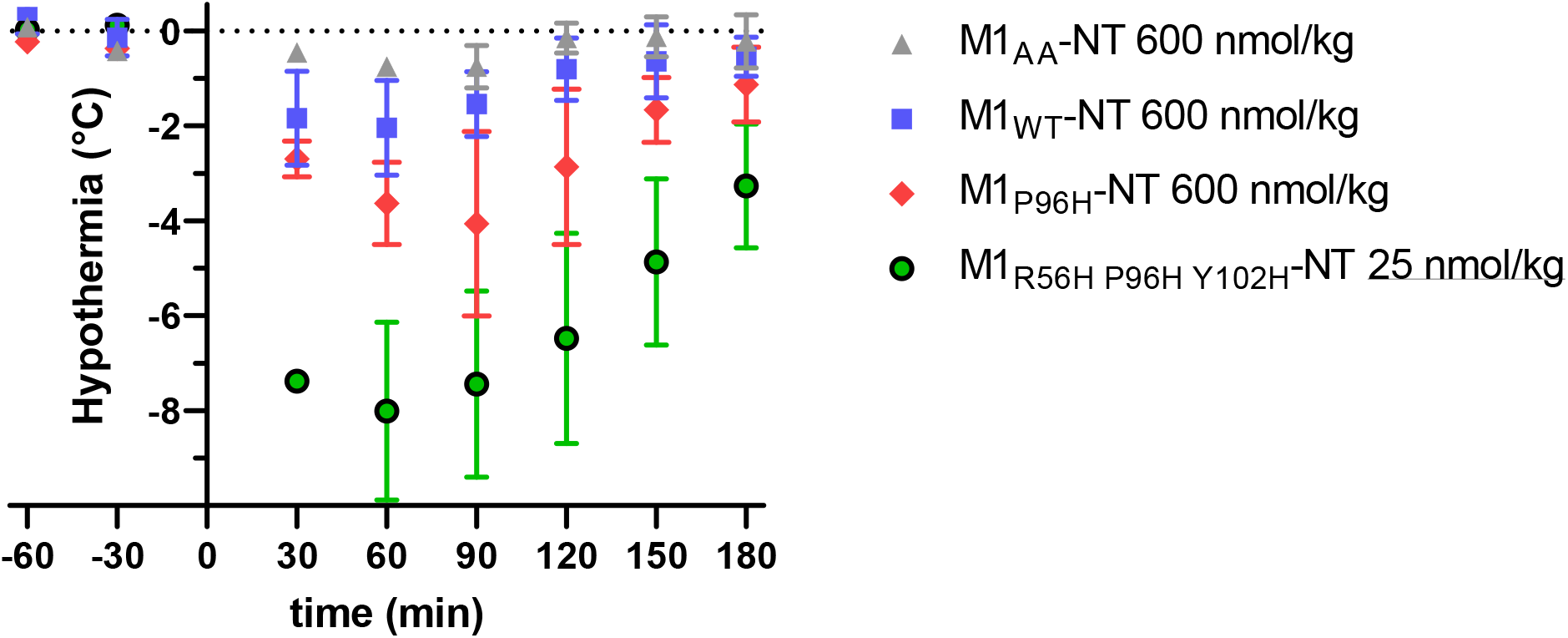
M1 mTfR binding nanobody–NT fusions for assessment of central nervous system target engagement. Blinded assessment of M1 variants injected iv into wild-type mice (n = 3 / group). M1 and M1_P96H_ mutant caused modest hypothermia at 600 nmol/kg. M1_AA_ (non TfR binding) TfR caused no detectible hypothermia. M1_R56H, P96H, Y102H_ caused pronounced hypothermia at 25 nmol/kg.

### Rapid shuttling of macromolecular cargoes across the BBB *in vivo*

As noted above, NT-induced hypothermia can be considered an efficient screening assay. However, it is not a definitive test of generalized BBB transcytosis. Therefore, we used two additional methods to assess the capacity of M1_R56H, P96H, Y102H_ to shuttle macromolecular cargoes across the BBB *in vivo*: capillary-depleted brain homogenates and *in situ* histochemistry. For proof-of-concept assessments, we created a three nanobody construct: M1_R56H, P96H, Y102H_ plus two tandem copies of the P2X7 receptor binding nanobody 13A7 [42] produced as a single protein using glycine-serine linkers between nanobodies (**Figure 3A**). As a non-binding control, we created a construct with M1_R56H, P96H, Y102H_ and two tandem copies of the human amyloid-beta receptor binding nanobody Nb3 [29], [43] (**Figure 3B**). Nb3 has no known targets in the wild-type mouse brain. The three nanobody constructs were produced at final levels of 1 – 10 mg per 100 mL of culture media in *E. coli.* We biotinylated the three nanobody constructs at random lysines to facilitate detection. Biotinylation at 5:1 stoichiometry yielded an average of 1.05±0.1 to 1.5±0.06 biotin adducts per nanobody, indicating relatively sparse labeling given that there are 4-5 lysines per nanobody.

**Figure 3.**
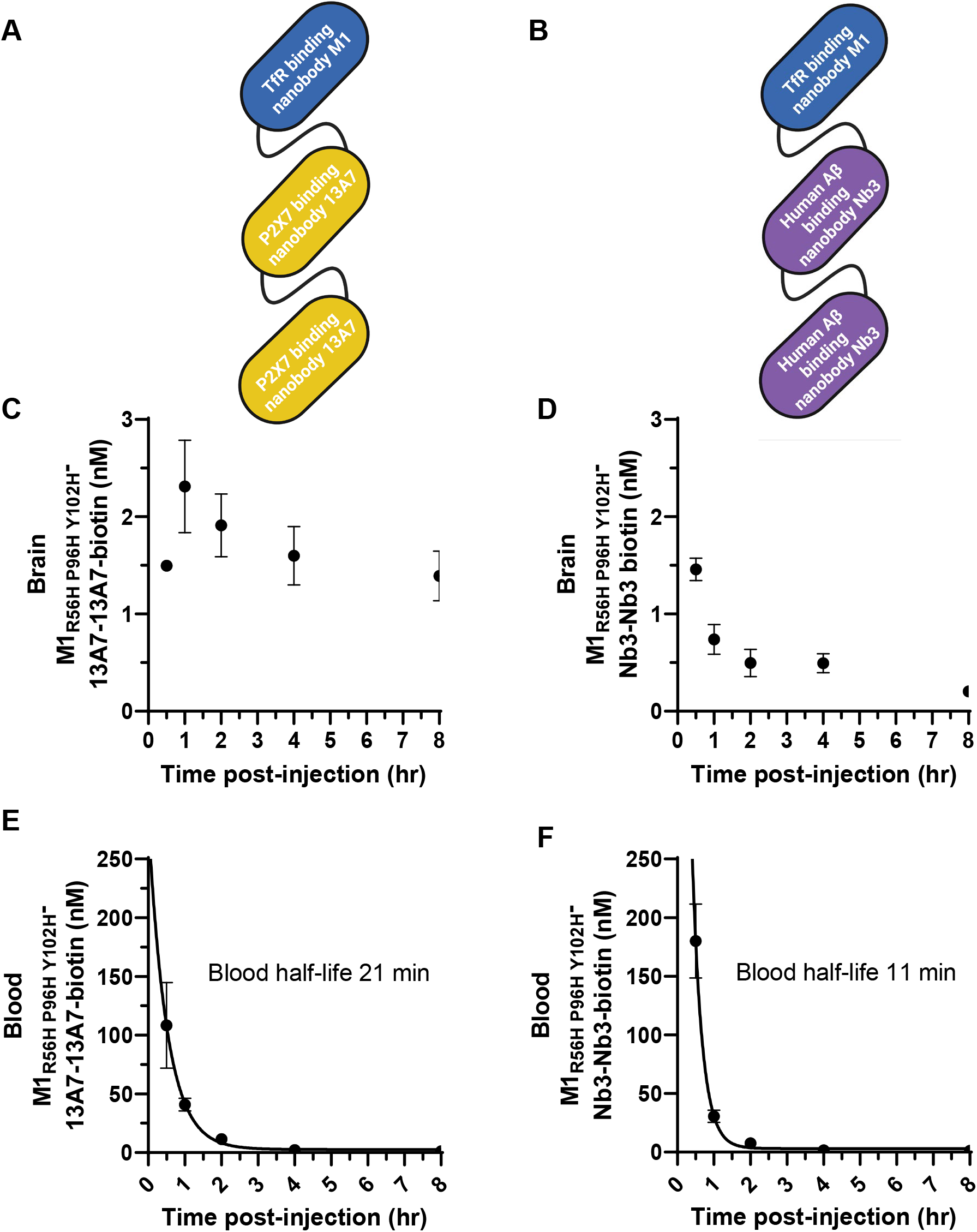
BBB transcytosis of macromolecular cargos by M1_R56H, P96H, Y102H_ and brain target engagement *in vivo*. **A.** Diagram of the three nanobody construct including M1_R56H, P96H, Y102H_ plus a tandem dimer of the P2X7 receptor binding nanobody 13A7 for brain target engagement. **B.** Diagram of a control three nanobody construct including M1_R56H, P96H, Y102H_ plus a tandem dimer of the human amyloid-beta binding nanobody Nb3 which has no known targets in the mouse brain. **C.** Time course of capillary depleted lysate levels of biotinylated M1_R56H, P96H, Y102H_ -13A7-13A7 (n=3 per time point) after iv injection of 30 nmol/kg. **D.** Time course of capillary depleted lysate levels of biotinylated M1_R56H, P96H, Y102H_ -Nb3-Nb3 (n=3 per time point) after iv injection of 30 nmol/kg. **E-F.** Time course of blood levels of the biotinylated three nanobody constructs from the same mice as panels C-D.

There was clear evidence of BBB transcytosis of the M1_R56H, P96H, Y102H_ 13A7-13A7 construct after intravenous injection in mice. The biotinylated three nanobody construct was readily detected in capillary-depleted brain homogenates after iv injection of 30 nmol/kg. Capillary-depleted lysate levels rose from 30 minutes to 1 hour, peaked at 1 hour, and remained approximately 60% of peak at 8 hours (**Figure 3C**). Levels of the control BBB crossing three nanobody construct in which 13A7 was replaced by Nb3 followed different kinetics, with similar levels at 30 minutes but <15% retained at 8 hours (**Figure 3D**). At 8 hours, the brain levels of M1_R56H, P96H, Y102H_ 13A7-13A7 were ∼7 fold higher than M1_R56H, P96H, Y102H_-Nb3-Nb3 (1.39 vs. 0.2 nM). Two way ANOVA was used to analyze the capillary depleted brain homogenate data. There were significant main effects of both construct (M1_R56H, P96H, Y102H_ -13A7-13A7 vs. M1_R56H, P96H, Y102H_ -Nb3-Nb3; F_1,20_ =151, p<0.0001) and time

(F_4,20_ =9.9, p=0.0001), as a well as a significant construct x time interaction (F_4,20_ =9.7, p=0.0002). Post-hoc comparisons were significant for all time points except 30 minutes (p<0.0001). Capillary depletion was approximately 98-99% effective, based on Claudin 5 ELISA measurements in crude vs capillary depleted lysates (**Suppl. Fig. 2A**). Western blots for VE cadherin confirmed capillary depletion (**Suppl. Fig. 2B**). Biotinylated nanobody construct levels in the capillary fractions were elevated at 30 minutes and 1 hour but below the limit of reliable quantitation (<0.166 to 0.2 nM) at later times (**Suppl. Fig. 3A-B**). The time course of the constructs in blood showed a mono-exponential decline with half-lives of approximately 21 and 11 minutes respectively (**Figure 3E-F**). These capillary depleted lysate results were compatible with the hypothesis that M1_R56H, P96H, Y102H_ was capable of shuttling proof-of-concept biological macromolecules across the BBB *in vivo* and that brain kinetics depended on brain target engagement.

### Prolonged blood half-life increases peak brain levels

In order to determine whether higher brain concentrations of nanobody constructs could be obtained by prolonging blood half-life, we added an albumin binding nanobody called Nb80 [44] making a four nanobody construct (**Figure 4A**). TfR binding, P2X7 receptor binding, and albumin binding functions were retained in the biotinylated four nanobody constructs, with characteristics comparable to those of the individual nanobodies. A control four nanobody construct in which 13A7 was replaced with Nb3 was also produced. The four nanobody constructs were produced at final levels of 1 – 10 mg per 100 ml of culture media in *E. coli.* The time course of the four nanobody construct in blood showed a mono-exponential decline with half-life of approximately 2.6 hours (**Figure 4B**).

**Figure 4:**
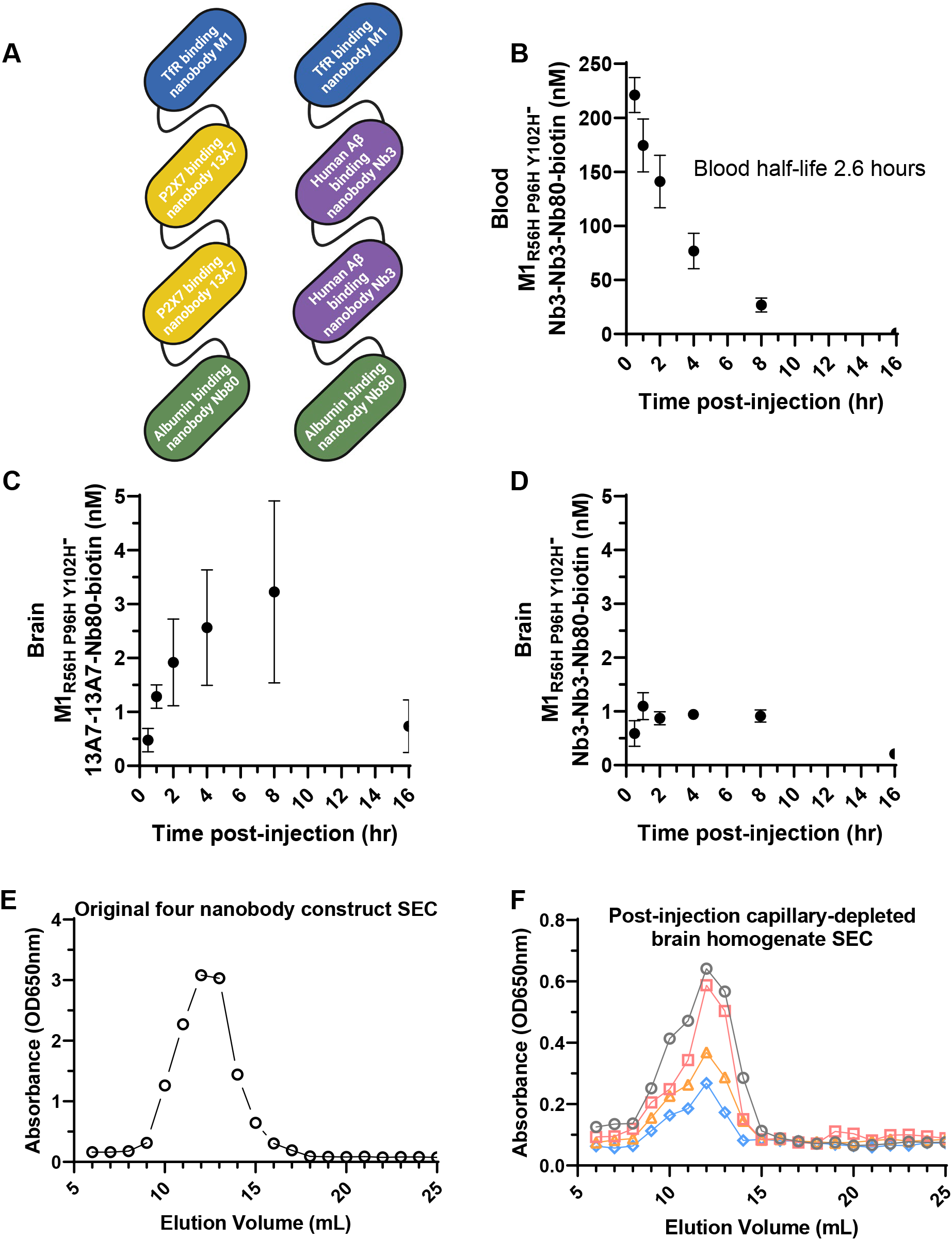
Blood half-life extension using an albumin-binding nanobody further enhances *in vivo* BBB transcytosis of macromolecular cargos by M1_R56H, P96H, Y102H_. **A.** Diagram of four nanobody constructs including M1_R56H, P96H, Y102H_, a tandem dimer of the P2X7 receptor binding nanobody 13A7 for brain target engagement or the human amyloid-beta binding nanobody Nb3 with no known mouse brain targets, plus the albumin binding nanobody Nb80. **B**. Time course of blood levels of the biotinylated four nanobody constructs demonstrating extended 2.6 hour half-life. **C.** Time course of capillary depleted lysate levels of biotinylated M1_R56H, P96H, Y102H_ -13A7-13A7-Nb80 (n=3 per time point) after iv injection of 30 nmol/kg. **D.** Time course of capillary depleted lysate levels of biotinylated M1_R56H, P96H, Y102H_ -Nb3-Nb3-Nb80 (n=3 per time point) after iv injection of 30 nmol/kg. **E-F.** Size exclusion chromatography of a biotinylated M1_P96H_-13A7-13A7-Nb80 prior to injection (**E**) and in brain homogenates (**F**) from 4 different mice (gray, red, orange, and blue symbols) 2 hours after iv injection into wild-type mice at 300 nmol/kg body weight demonstrating integrity of the four nanobody constructs in brain. No substantial aggregation or degradation was apparent.

Again, BBB transcytosis of the four nanobody construct was clearly observed in capillary-depleted brain homogenates, with peak levels occurring later than for the three nanobody construct. Capillary-depleted lysate levels rose from 30 minutes to 4 hours, peaked at 4-8 hours at levels approximately 50% higher than the peak concentration attained by the three nanobody construct, and were >80% cleared at 16 hours (**Figure 4C**). In a two way ANOVA, there was a significant interaction between construct (M1_R56H, P96H, Y102H_ -13A7-13A7-Nb80 vs. M1_R56H, P96H, Y102H_ -13A7-13A7) and time (F_1,20_ =4.5, p=0.0094). Post-hoc comparisons were only significant for the 8 hour time point (p=0.027); the differences at earlier time points were not statistically significant after correction for multiple comparisons. Levels of an otherwise identical four nanobody construct including the M1_AA_ mutation that does not bind TfR nor cross the BBB were undetectable. Levels of a BBB crossing four nanobody construct in which 13A7 was replaced by Nb3, the anti-human amyloid-beta nanobody that does not bind any known targets in the wild-type mouse brain, peaked at approximately 3 fold lower levels than the P2X7 receptor binding construct (**Figure 4D**). Two way ANOVA was again used to analyze the capillary depleted brain homogenate data. There were significant main effects of both construct (M1_R56H, P96H, Y102H_ -13A7-13A7-Nb80 vs. M1_R56H, P96H, Y102H_ -Nb3-Nb3-Nb80; F_1,24_ =18.2, p=0.0003) and time (F_5,24_ =5.8, p=0.0012), as well as a significant construct x time interaction (F_5,24_ =2.95, p=0.03). Post-hoc comparisons were significant for the four hour (p=0.034) and eight hour (p=0.0014) time points. Biotinylated four nanobody construct levels in the capillary fractions were elevated at 30 minutes and 1 hour, but below the limit of reliable quantitation (<0.2 nM) at later times (**Suppl Fig. 3C-D**). Four nanobody constructs remained intact and were not detectibly degraded in the capillary depleted brain lysates; size exclusion chromatography revealed similar distribution of sizes in the capillary depleted lysates compared to the original injected material (**Figure 4E-F**). These capillary depleted lysate results were compatible with the hypothesis that a modest prolongation of serum half-life of the M1_R56H, P96H, Y102H_ -based nanobody constructs produced higher peak brain levels of the biological macromolecule cargos and longer residence time in the brain.

Further confirmation of BBB crossing and brain target engagement was provided by *in situ* histochemistry using slices from the contralateral hemispheres of the same mice injected with four nanobody constructs described above. At 30 minutes after injection, the biotinylated four nanobody construct was detected in structures with morphologies consistent with capillaries (**Figure 5A**). At later times, cellular structures were labeled with increasing intensity (**Figure 5B-F**) in a pattern consistent with the *ex vivo* binding of the 13A7 nanobody construct (**Figure 5G**). Peak binding appeared at 4-8 hours, with reduced binding at 16 hours. The time course of *in situ* binding was similar to that observed in the capillary depleted brain lysates. There was no detectible *in situ* histochemical signal 4 hours after injection of the construct containing the M1_AA_ mutation that does not bind TfR nor cross the BBB (**Figure 5H**). There was faint capillary-like staining but no detectible cellular structures on *in situ* histochemical signal after injection of the four nanobody construct in which 13A7 was replaced by Nb3 (**Figure 5I**). Additional histochemical images at lower magnification including other brain regions are shown in **Suppl. Fig. 4**. These *in situ* histochemistry results provided further support for the hypothesis that M1_R56H, P96H, Y102H_ was capable of shuttling proof-of-concept biological macromolecules across the BBB *in vivo* where they could specifically interact with brain targets.

**Figure 5:**
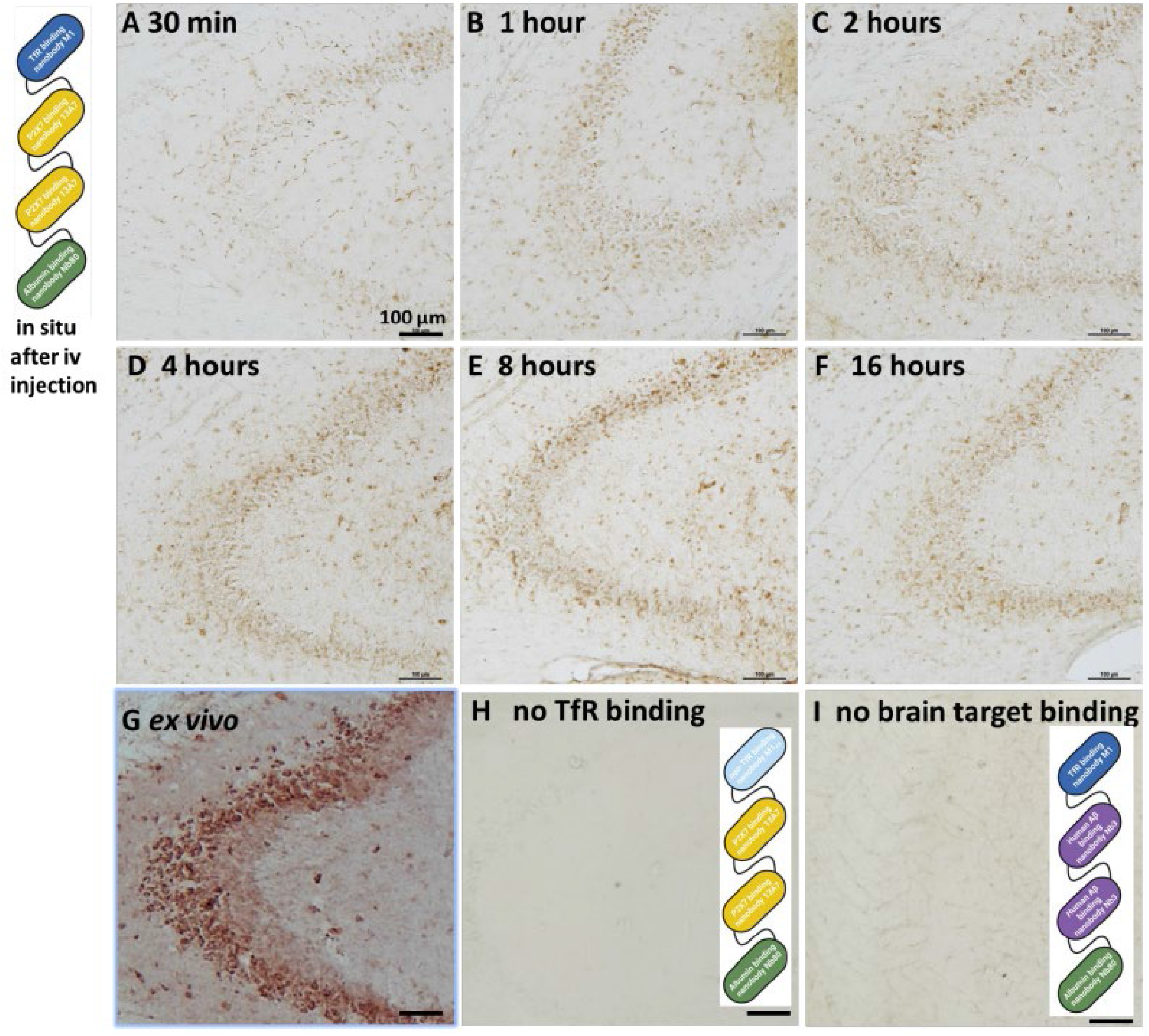
I*n vivo* target engagement of four nanobody constructs after iv injection. **A-F.** *In situ* labeling of biotinylated M1_R56H, P96H, Y102H_-13A7-13A7-Nb80 injected iv into wild-type mice at 600 nmol/kg body weight at 30 min showing largely capillary labelling, and at 1, 2, 4, 8 and 16 hrs showing cellular labeling in hippocampus. **G.** Positive control: *ex vivo* staining of naïve mouse brain slice with 50 nM biotinylated M1_R56H, P96H, Y102H_-13A7-13A7-Nb80. **H.** Negative control: no *in situ* labeling after intravenous injection of 600 nmol/kg M1_AA_ -13A7-13A7-Nb80 which does not bind mTfR. **I.** Negative control: trace capillary *in situ* labeling after intravenous injection of 600 nmol/kg M1_R56H, P96H, Y102H_ -Nb3-Nb3-Nb80. Nb3 has no known targets in wild-type mouse brain.

### Nonlinear relationship between administered dose and brain levels of shuttled macromolecular cargo

The utility of a BBB shuttle system for specific diagnostic and therapeutic applications may depend on the quantitative relationship between administered dose and achievable brain levels. We assessed the capillary depleted brain homogenate levels 4 hours after injection of a wide range of doses of the biotinylated TfR and P2X7 receptor binding four nanobody construct (**Figure 6**). As has been reported for another TfR-binding antibody fragment [45], a strong saturation effect was observed; peak capillary depleted brain homogenate levels were observed at 30 nmol/kg (approximately 1.8 mg/kg), with no increased levels after injections of much higher doses. At the lowest doses administered, 1 and 3 nmol/kg, the brain levels of 0.64 and 1.21 nM indicated that the construct was transported across the BBB at 13.4 and 8.8% of the injected dose/gram of brain tissue. At 30 nmol/kg, levels as high as 4.5 nM indicated that the construct was transported across the BBB at 3.5% injected dose/gram of brain tissue. Full calculations are shown in **Suppl. Table 1**. There were small *decreases* in capillary depleted brain homogenate levels after injection of higher doses. We performed mathematical curve fitting of two models to the dose response data. The first model involved fits to the standard Hill equation with 3 parameters, and the alternative model involved fits to a 3 parameter model that included substrate inhibition. The alternative model fit the data better (r^2^ = 0.75 vs. 0.69). These results are compatible with the hypothesis that M1_R56H, P96H, Y102H_ can shuttle proof-of-concept biological macromolecules across the BBB efficiently at low to moderate doses but becomes substantially less efficient at higher doses due to both saturation and an apparent substrate inhibitory effect.

**Figure 6:**
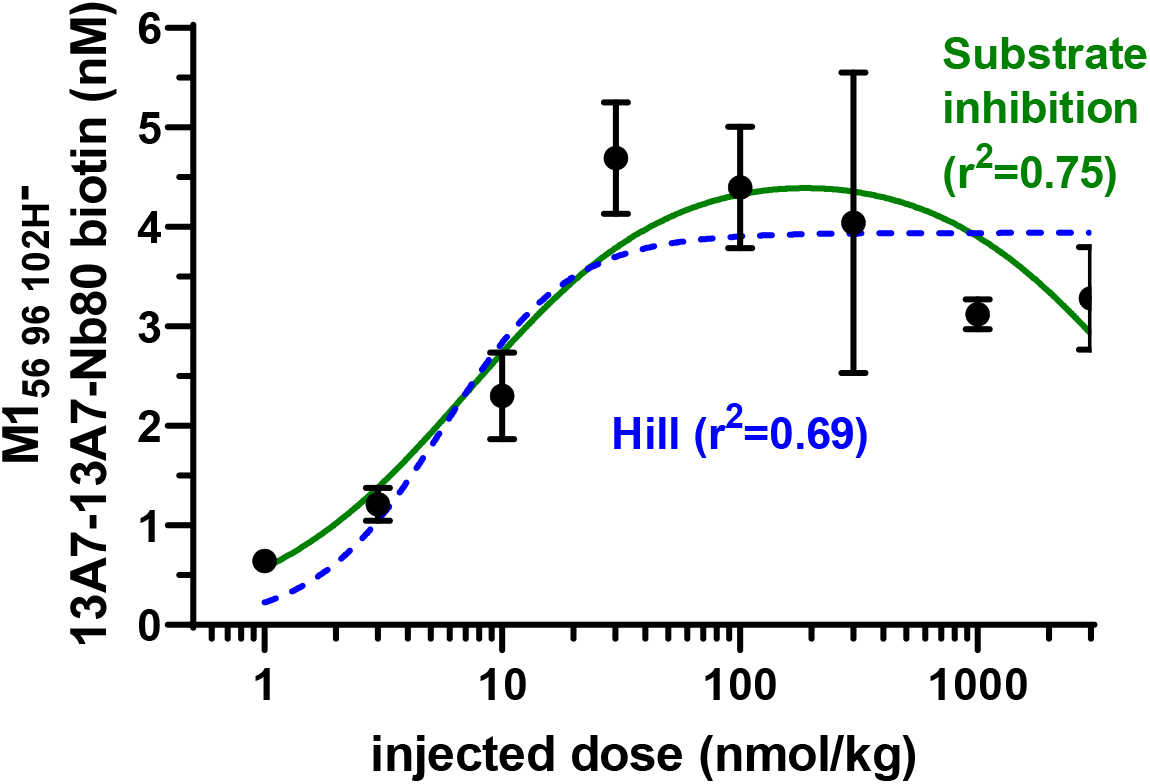
Capillary depleted brain lysate levels of nanobody construct 4 hours after injection of a range of doses of biotinylated M1_R56H, P96H, Y102H-_13A7-13A70-Nb80 in mice. N=3 per dose, except for N=2 at 1 nmol/kg and N=2 for 1000 nmol/kg. Hill equation fit parameters were B_max_=3.94 nM, ED_50_=5.6 nmol/kg, Hill coefficient=1.6. Substrate inhibition fit parameters were B_max_=4.7 nM, ED_50_=7.3 nmol/kg, K_i_=4877 nmol/kg.

## DISCUSSION

In summary, the mouse TfR-binding nanobody M1 with three histidine mutations can be used as a shuttle to transport other nanobodies across the intact BBB in mice. After intravenous injections of low to moderate doses, 1-30 nmol/kg, nanobody construct levels in capillary depleted brain lysates reached over 3.5% of injected dose per gram of brain, which is comparable to other top performing receptor mediated transcytosis shuttle systems (**Suppl Table 2**). The kinetics of M1 nanobody-mediated BBB transcytosis were relatively fast. For a three nanobody construct, peak levels occurred at 1 hour, and for a four nanobody albumin binding construct peak levers were noted at 4-8 hours after injection with ∼80% clearance from the brain by 16 hours. The pH dependence of M1 histidine mutant binding partially correlated with *in vivo* BBB transport efficacy. This result is compatible with the hypothesis that unbinding of M1-based constructs in the relatively acidic environment of the intracellular vesicular compartments involved in transcytosis may favor fast delivery across the BBB (**Figure 7**).

**Figure 7:**
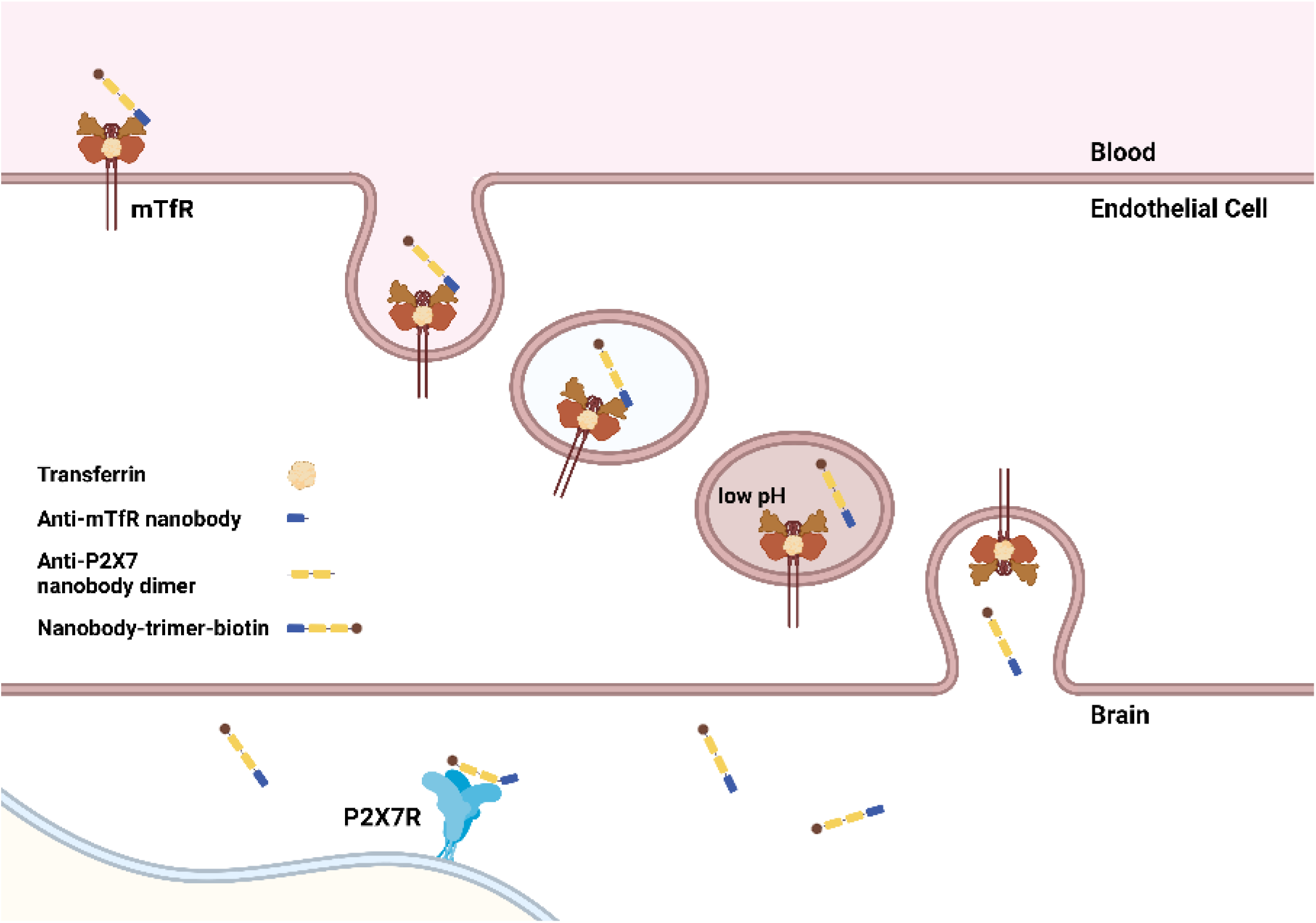
Diagram of hypothesized mechanisms involved in BBB transcytosis of mTfR-and-P2X7-receptor binding nanobody constructs. Anti-mTfR nanobodies bind to mTfR in blood and constructs are endocytosed. Nanobody constructs unbind from mTfR in a pH-dependent fashion in intracellular vesicular compartments. Nanobody constructs are released into the brain and bind P2X7 receptor targets. (Produced using BioRender)

It is likely that there will not be a “one size fits all” best solution to the problem of transporting macromolecules across the BBB. Nanobody-based receptor mediated transcytosis shuttles such as the pH-dependent transferrin receptor binding system presented here may play an important context-specific role. Nanobodies can be readily engineered, inexpensively produced in bacterial and yeast systems, and stored at room temperature. Furthermore, the BBB transport kinetics were relatively fast, especially for the 3 nanobody constructs. Thus, applications requiring frequent re-engineering, low cost, simplified logistics, and rapid kinetics may be favored. Examples could include development of a family of modular molecular imaging agents created by swapping out the P2X7 receptor binding nanobodies for other diagnostic marker-specific recognition domains. The relatively fast kinetics are potentially compatible with the workflow of diagnostic imaging involving injection of a contrast agent and scanning a few hours later. We have found that these nanobody constructs could be biotinylated via lysine conjugation without loss of efficacy, making it likely that conjugation of other labels such as metal chelators and fluorophores should also be feasible. A similar approach could be used to develop a family of relatively fast acting therapeutic agents, again by swapping out the P2X7 receptor binding nanobodies for other disease-specific therapeutic payloads. Of note, the first nanobody-based therapeutic approved for human use, caplacizumab for TTP, has rapid kinetics appropriate for acute treatment and once daily dosing [46].

The nanomolar levels of the M1 nanobody-based constructs detected in capillary depleted brain lysates after intravenous injection of relatively modest doses are potentially compatible with many applications. PET scans using radioactive metal conjugates may be feasible given the high percent injected dose/gram of brain and high ratio of concentrations for active constructs vs non-target binding controls. Therapeutic applications should also be feasible for agents with relatively high affinity. It remains to be determined whether molecular contrast MRI using extremely small iron oxide nanoparticles [47] or other MRI contrast agents will be feasible. Fundamentally, the major limitation to applications requiring higher brain concentrations is the saturation of the transferrin receptor-based transport system. At injected doses above 30 nmol/kg no further increases in concentrations in capillary depleted brain lysates were observed. Saturation of the transferrin receptor-based transport system has been observed by others as well [45] (**Suppl. Table 2**), and to our knowledge transport of macromolecules via any receptor mediated transport system has not reached levels above ∼13 nM in capillary depleted brain lysates [31]. The relationship between capillary depleted brain lysate concentrations and brain extracellular space concentrations has not been determined (see Supplemental Discussion).

The 13A7 P2X7 receptor binding nanobody was chosen primarily for demonstration of proof-of-concept, but BBB penetrating P2X7 receptor binding agents have many potential therapeutic applications. Brain P2X7 receptor activation has been implicated in the pathophysiology of traumatic brain injury, post-traumatic headache, ischemic and hemorrhagic stroke, multiple sclerosis, neurodegenerative disorders, depression, and other conditions [48] [49] [50] [51] [52] [53] [54] [55] [56] [57] [58]. Development of P2X7 receptor binding reagents has been an active area of pharmaceutical research, and there remains an unmet need for highly specific, brain-penetrating P2X7 receptor antagonists [59]. Administration of a P2X7 receptor binding nanobody intracerebroventricularly has been shown to reduce stroke lesions in a mouse model but P2X7 receptor binding nanobodies were not effective after intravenous administration [56], further emphasizing the importance of BBB penetration. However, the specificity of the 13A7 nanobody has not been fully established, and alternative nanobodies that bind to human P2X7 receptors may be more appropriate for human therapeutic use [42] [56] [60].

There are several limitations to these findings. First, our strategy for introducing histidine mutations to impart pH dependent binding was empirical rather than structure based. Despite intensive efforts, we and other have not yet been able to obtain high resolution structural information on the mouse TfR extracellular domain. If mouse TfR structures can be obtained, it would open the doors for further rational mutagenesis to impart even sharper pH dependent binding which in turn could further improve transcytosis. The structure of the human TfR extracellular domain has been solved [61] [62], so rational mutagenesis could be used to accelerate development of pH-dependent human TfR-binding nanobodies. Second, we have not yet been able to directly measure nanobody on and off rates at pH 5.5 and pH 7.4 for technical reasons; our measurements of pH-dependent apparent unbinding may not reflect true kinetics. Third, we have not performed detailed tracking of the transcytosis of the M1-based nanobody constructs. We do not know exactly where the presumed pH-dependent unbinding occurs from a cell biological perspective. Likewise, we do not understand the mechanisms of clearance from the brain nor whether these clearance mechanisms are pH-dependent. Fourth, we have not performed a full quantitative assessment of the relationships between *in vivo* NT-induced hypothermia effects and *in vivo* concentrations in capillary depleted brain lysates for all of the tested constructs. We performed *in vivo* measurements in capillary depleted brain lysates only for our lead candidate, the M1_R56H, P96H, Y102H_ mutant with the largest NT-induced hypothermia effect. NT-induced hypothermia likely reflects transcytosis across the BBB in the hypothalamus [63], and the hypothalamic BBB may not be identical to that of other parts of the brain, so the relationship between *in vivo* assays is not necessarily 1:1. Fifth, while we have not observed overt toxicity after single dose injection of M1 nanobody constructs apart from one mouse that died shortly after injection, formal toxicity testing after single dose and multidose injection has not yet been performed. Sixth, we have not assessed the BBB transport capacity of the M1 nanobody mutants for other macromolecular cargoes such as full size antibodies, single chain variable fragments, enzymes, or nucleic acids. Relatedly, it is not clear whether brain cytosolic targets can be reached even when BBB crossing is efficient; both the NT receptor and P2X7 receptor are extracellular, as are most of the targets of macromolecular cargos assessed in previous work in the field (**Suppl. Table 2**). Seventh, the M1 nanobody mutants bind to mouse but not human TfR. Additional nanobodies binding to human TfR are in development and we plan to use the same histidine mutagenesis strategy to impart similar pH-dependent binding to human TfR binding nanobodies. There are other examples of human specific TfR binding nanobodies that could also be engineered in this fashion [27]. Likewise, we have not yet ‘humanized’ our nanobody constructs, but the framework for doing so has been relatively well established [64]. It will be most appropriate to humanize nanobody constructs that bind to human targets. Finally, we have not yet used the M1 nanobody constructs for preclinical imaging or therapeutic studies. Imaging studies using the 3 nanobody constructs and fast-acting therapeutic studies using the 4 nanobody constructs are the logical next steps in this line of research given the rapid kinetics of these constructs and the important role of P2X7 receptors in many disease processes.

### CONCLUSIONS

In conclusion, our results are compatible with the hypothesis that an engineered TfR-binding nanobody with pH-dependent unbinding has potential to serve as a shuttle system for transcytosis of biological macromolecules across the BBB and into the brain. Additional development will be required to determine whether this or other nanobody-based shuttle systems will be useful for diagnostic and therapeutic applications.

## LIST OF ABBREVIATIONS

BBB: blood brain barrier
BSA: bovine serum albumin
HABA: 4’-hydroxyazobenzene-2-carboxylic acid
HBSS: HEPES-buffered saline
IPTG: isopropylthio-β-galactoside iv: intravenous
NT: neurotensin
PBS: phosphate-buffered saline
RIPA: radioimmunoprecipitation assay
scFv: single chain variable fragment
SEC: size exclusion chromatography
TfR: transferrin receptor
WT: Wild-type

## Acknowledgements

The authors would like to thank Ms. Kathy Ireland, Dr. Arthur Kellermann, Dr. Dale Kiesewetter, Dr. Alan Koretsky, Dr. Walter Koroshetz, Dr. Lorna Role, Ms. Kathy Scherer and Dr. Nina Schor for their support, and multiple colleagues for stimulating discussions.

## Author Contributions

DLB and TJE designed experiments. TJE, CF, ER, JHK, and SS performed experiments. DLB, TJE, and CF analyzed data. DLB, TJE, and JHK produced figures. DLB wrote the first draft of the manuscript, with revision and contributions from all authors.

## Data and material availability

All protein and DNA sequences will be made available by request from the authors. Recombinant purified nanobody constructs and DNA vectors are available for sharing under appropriate material transfer agreements.

## Declarations

### Ethics approval

All animal experiments were conducted according to protocols approved by the NINDS intramural research program animal studies committee, protocol # 1406

### Consent for publication

Clearance has been obtained from the NINDS intramural research program and from the Uniformed Services University

### Disclaimer

The views, information or content, and conclusions presented do not necessarily represent the official position or policy, nor should any official endorsement be inferred, on the part of the National Institutes of Health, the Uniformed Services University, the Department of Defense, Henry M. Jackson Foundation for the Advancement of Military Medicine, Inc., or other government agency.

## SUPPLEMENTAL INFORMATION

### Supplemental Methods

#### Quantification of nanobody biotinylation

The degree of biotin incorporation was determined using the Pierce Biotin Quantitation Kit (#28005, Thermo Scientific). The assay measures the displacement of avidin from HABA (4’-hydroxyazobenzene-2-carboxylic acid) with nanobody-conjugated biotin. Briefly, the HABA/avidin complex was reconstituted in ultra-pure water and a 1/10 dilution was prepared in 1xPBS. The absorbance at 500nm was recorded for a baseline measurement. The biotinylated nanobody samples were subsequently diluted in the HABA/avidin reagent, incubated briefly and the absorbances at 500nm were measured. The Δ500nm measurement of the reagent alone and diluted biotin-nanobody were then used to determine the molar equivalents of biotin in the nanobody sample as outlined in the manufacturer’s operating protocol.

#### Quantification of Claudin-5 by ELISA

The concentration of claudin-5 in capillary-depleted brain homogenates and the isolated capillary fractions were measured using a commercially available ELISA kit (#CSB-EL005507MO, Cusabio). Briefly, capillary-depleted homogenates and isolated capillary fractions were prepared and supplemented with 0.5% (v/v) radioimmunoprecipitation assay (RIPA) buffer. Total protein was measured for each sample using a micro-BCA protein assay (#23235, Thermo Scientific). The claudin-5 assay was performed according to the manufacturer’s protocol, with the minor modification of supplementing the standard curve and all dilutions with RIPA buffer at 0.5% (v/v) to normalize any effect of RIPA buffer on assay binding. The samples and standard were loaded onto the ELISA plate and incubated overnight at 4°C. The assay was then developed following the manufacturer protocol and the absorbance measured at 450nm. The sample concentrations were normalized to total protein as reported in picograms claudin-5/milligrams total protein.

#### VE-Cadherin Western blot in mouse brain homogenates

Naïve wild-type mouse brain homogenates were processed to deplete the capillary component as described above. Gel electrophoresis was performed by separating 20-micrograms of total protein per lane of both pre-capillary-depletion and post-capillary-depletion brain homogenates on a 10% Bis-Tris acrylamide gel, including size standards. Following electrophoresis, the gel was transferred to 0.2 µm nitrocellulose blotting membrane using the wet-transfer method in tris-glycine buffer at 100 volts for 2hr in a chilled transfer tank. The transferred membrane was blocked using a 3% (w/v) bovine serum albumin in 1xPBS solution for 1 hour with gentle agitation. The VE-cadherin primary antibody (#ab205336, Abcam) was diluted 1:1000 (v/v) in blocking buffer and exchanged onto the membrane and allowed to incubated overnight at 4°C with gentle agitation. Following three washes with 1xPBS for 10 minutes each, goat anti-rabbit-peroxidase antibody (#ab97051) solution was prepared in the blocking buffer and exchanged onto the membrane for 1 hour at room-temperature with gentle agitation. A final triple wash was performed, and the membrane was developed using Clarity Max Western ECL (#1705062, BioRad) reagent and the chemiluminescence imaged on a BioRad ChemiDoc image station using standard acquisition parameters.

#### Supplemental Results and Figures

We wished to assess the relative importance of affinity at pH 7.4 vs. pH dependent unbinding under acidic conditions for *in vivo* NT-mediated hypothermia. To do so, we plotted the extent of NT-induced hypothermia after intravenous injection of 25 nmol/kg of each M1-NT construct vs. each parameter (**Suppl. Fig. 1A-B**). We found that there was no significant monotonic relationship between affinity and NT effect (Spearman r=0.01, p=0.96); there were NT constructs with strong NT effects with affinities ranging from 6 to 246 nM, and other NT constructs with similar affinities that did not have strong NT effects. Instead, there was a clearer correlation of NT-induced hypothermia *in vivo* with the fold increase in unbinding at pH 5.5 vs. 7.4 (Spearman r=0.4, p=0.004). However, even this parameter only explained a modest proportion of the variance in NT-induced hypothermia. Notably, several NT constructs with substantially increased unbinding at pH 5.5 such as M1_R56H, P96H, Y102H_ still did not induce hypothermia *in vivo.*In a three-dimensional plot of affinity at pH 7.4 and pH-dependent unbinding vs. NT-induced hypothermia, we observed that the combination of affinity in the range of 6 to 246 nM plus at least 2-fold greater unbinding at pH 5.5 vs pH 7.4 was strongly associated with potent NT-induced hypothermia (**Suppl. Fig. 1C**). There were still outliers; the M1_Y98H, Y102H_ mutant for example mediated substantial NT-induced hypothermia (5°C), with only modestly greater unbinding at pH 5.5 vs pH 7.4 and a low affinity (>5000 nM). Thus, it appears that both optimal affinity and pH dependent unbinding contribute substantially but do not fully explain the NT-induced hypothermia mediated by the M1 histidine mutants. Nonetheless, the M1_R56H, P96H, Y102H_ mutant with strongly pH dependent unbinding and preserved high affinity was highly effective at producing NT-mediated hypothermia *in vivo*.

**Suppl. Fig.1.**
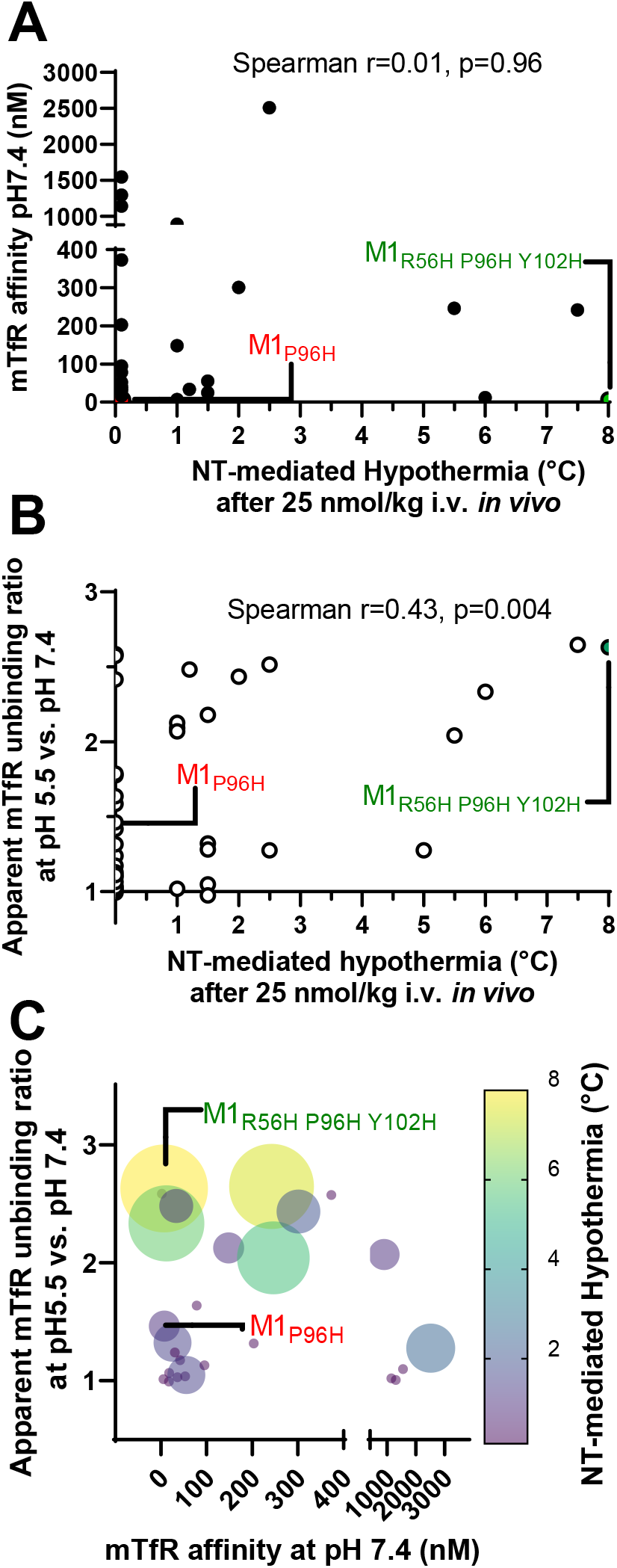
*In vitro* TfR binding vs. *in vivo* NT-mediated hypothermia for multiple M1 histidine mutants. **A.** NT-mediated hypothermia *in vivo* vs. mTfR binding measured by ELISA at pH 7.4 for constructs with measurable affinities. **B.** NT-mediated hypothermia *in vivo* vs. the ratio of apparent mTfR unbinding at pH 5.5 vs. unbinding at pH 7.4 as measured by ELISA. **C.** NT-mediated hypothermia vs. both affinity and apparent pH 5.5 vs pH 7.4 unbinding ratio. NT-mediated hypothermia expressed as both bubble size and color. NT-mediated hypothermia was measured in a blinded fashion after intravenous injection of 25 nmol/kg of each construct (n=3 mice per construct).

**Suppl. Fig. 2.**
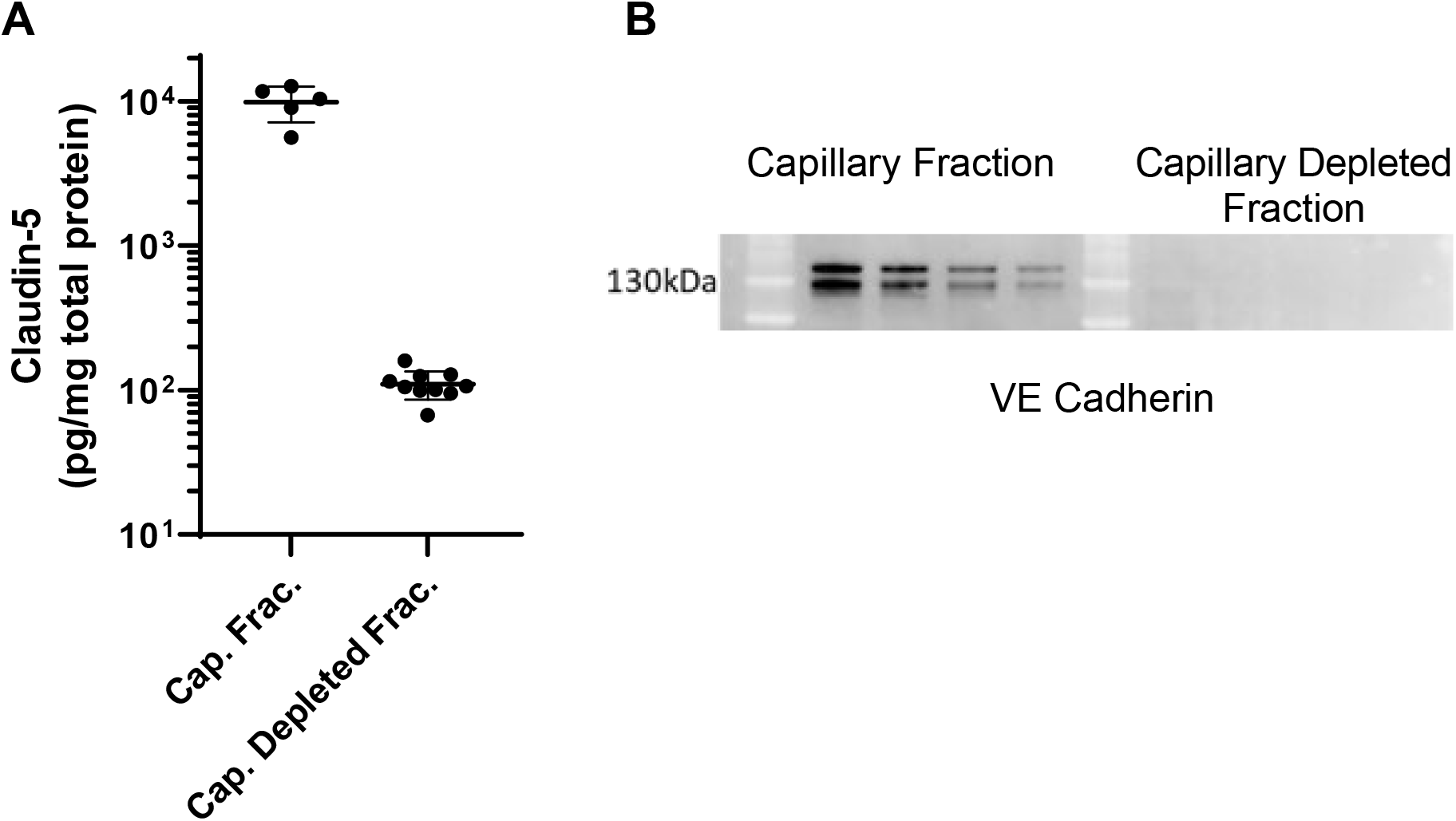
Verification of capillary depletion from brain lysates. **A.** Claudin-5 ELISA of lysates from the capillary containing fractions and capillary depleted fractions of mouse brains after injection of M1-based nanobody constructs. Note log scale of y-axis. Capillary depletion removes ∼99% of Claudin-5, a brain capillary endothelial marker. **B.** VE Cadherin western blotting of lysates from the capillary containing fractions and capillary depleted fractions. After capillary depletion, there was no detectible VE Cadherin, another brain capillary endothelial marker.

**Suppl. Fig. 3.**
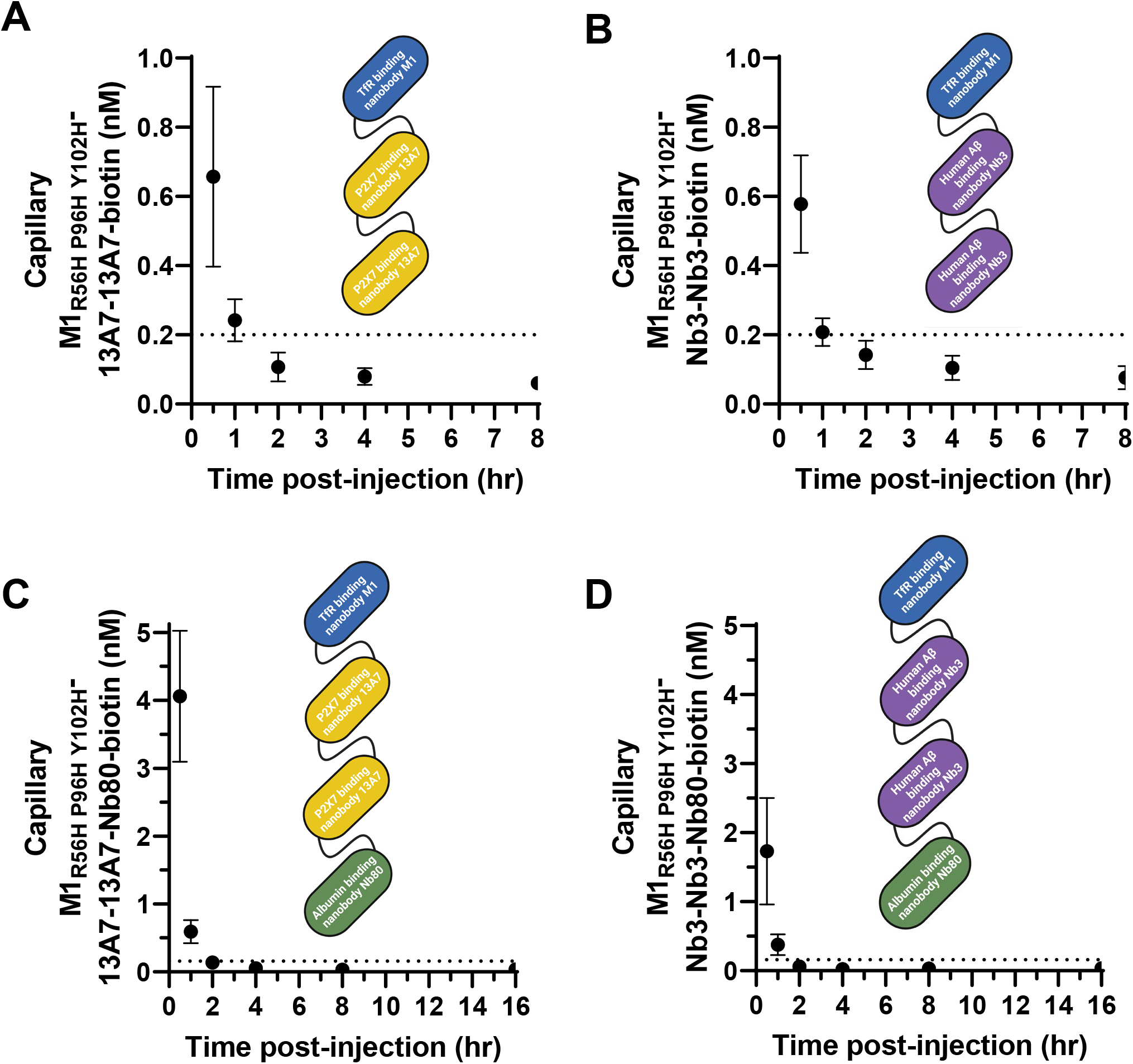
TfR-binding nanobody construct concentrations in capillary containing fractions of brain lysates. **A.** Three nanobody construct with P2X7 receptor target binding. **B.** Three nanobody construct with no brain target binding. **C.** Prolonged blood half-life four nanobody construct with P2X7 receptor target binding. **D.** Prolonged blood half-life four nanobody construct with no brain target binding.

**Suppl. Fig. 4.**
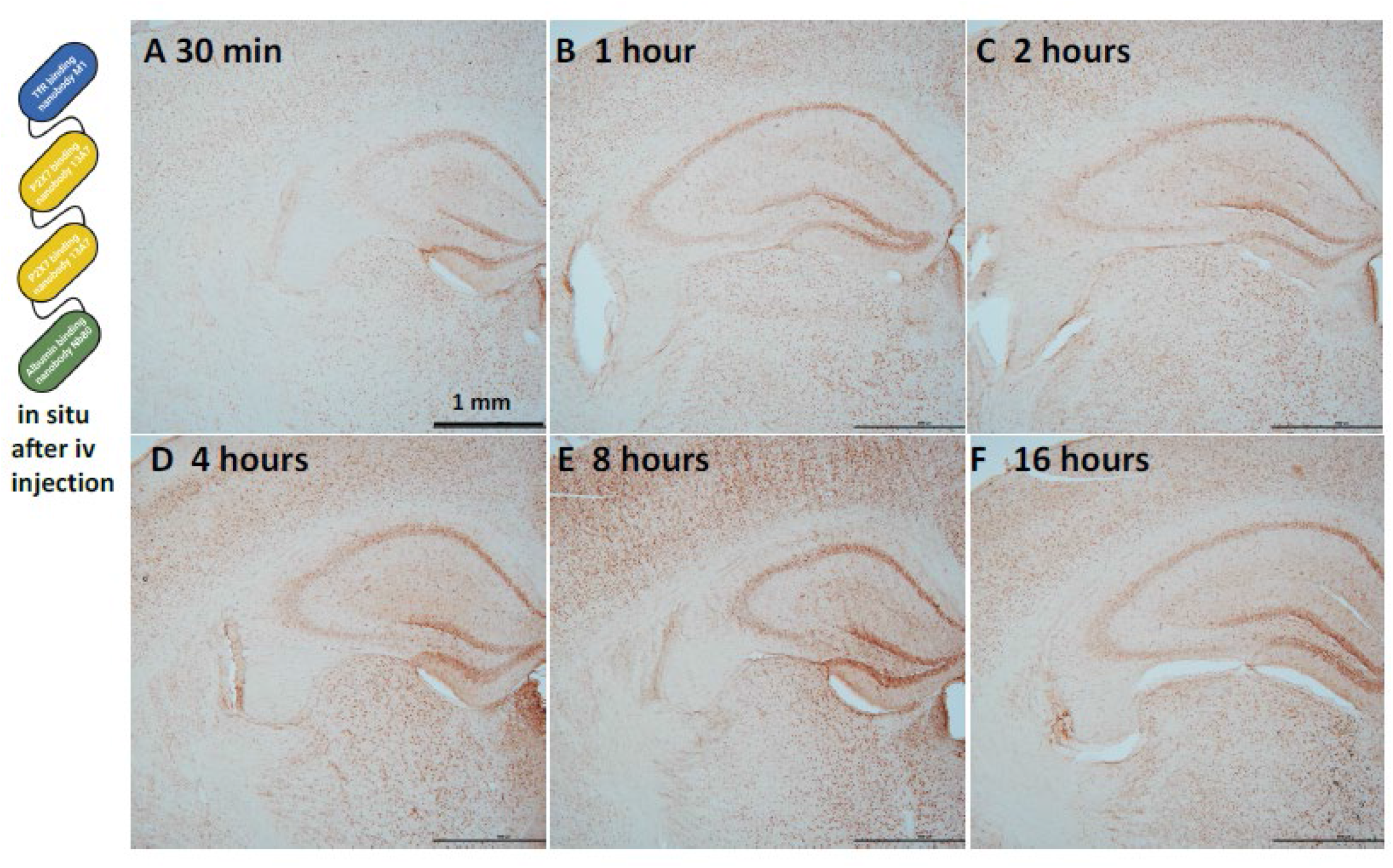
Lower magnification (4x) images from *in situ* labeling of biotinylated M1_R56H, P96H, Y102H_-13A7-13A7-Nb80 injected iv into wild-type mice at 600 nmol/kg body weight from the same mice as in Fig. 6. Both cortex and hippocampus at 30 min show capillary labelling. There was widespread cellular labeling at 1, 2, 4, 8 and 16 hrs in hippocampus, cortex, white matter and thalamus.

### Supplementary Tables

**Suppl. Table 1.**
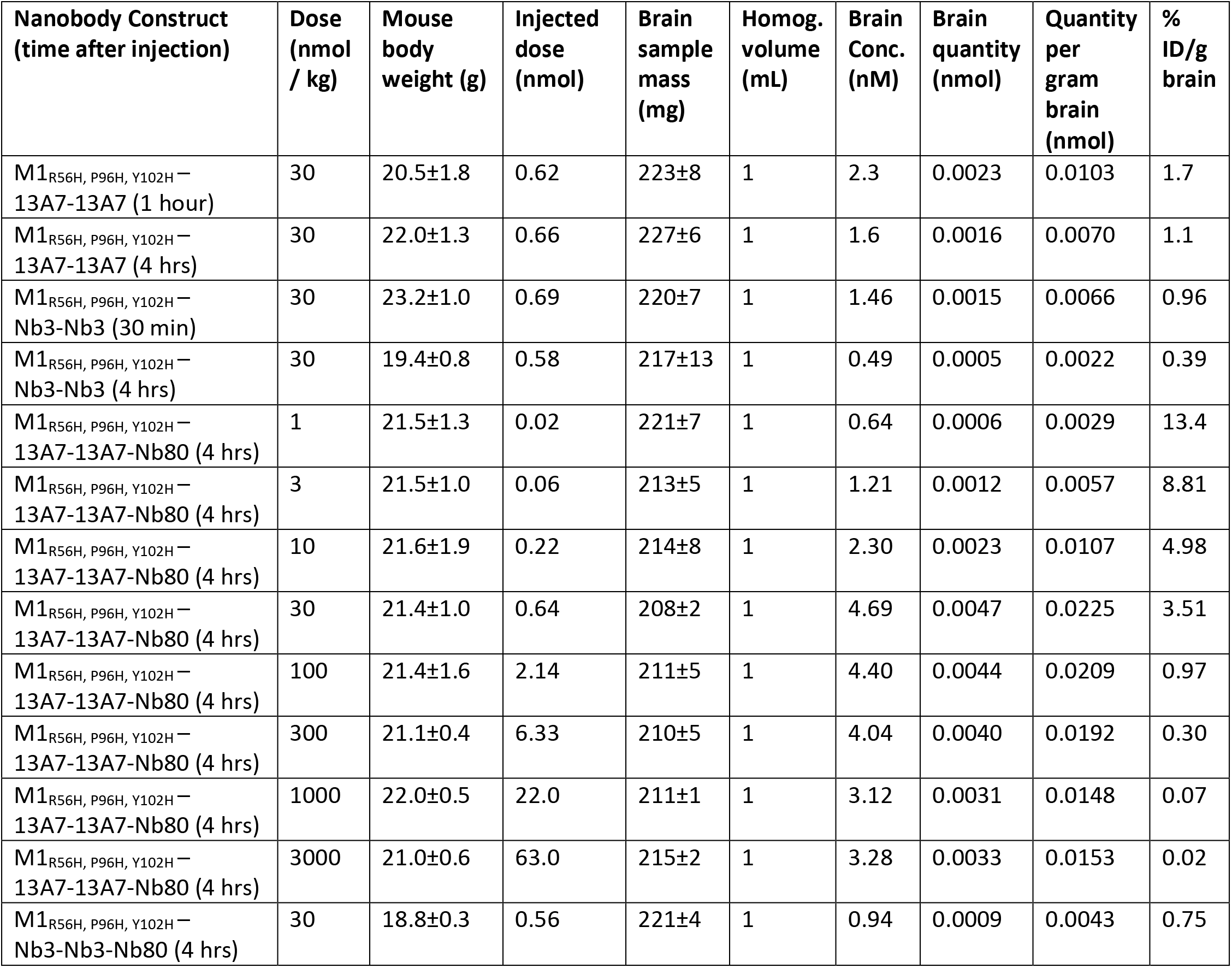
Percent injected dose per gram of brain (%ID/g) calculations (n=3 per group).

**Suppl. Table 2.** Percent injected dose/gram of brain for the P2X7 receptor binding M1_R56H, P96H, Y102H_-13A7-13A7 three nanobody construct and M1_R56H, P96H, Y102H_-13A7-13A7-Nb80 four nanobody construct compared with previously published BBB shuttles assessed using similar capillary depletion brain homogenate methods. When detailed methods were not specified, we assumed that mouse body weight was 20 grams and that 200 mg brain samples were lysed in 1 ml for these calculations. WT: Wild-type.

| Reference | BBB crossing approach | Macromolecule cargo | Dose injected | Time after injection | % Inj. Dose / gram Brain |
| --- | --- | --- | --- | --- | --- |
| Friden et al., 1991 [3] | Rat TfR IgG OX-26 | Methotrexate via Fc linked carbohydrate in rats | ~30 nmol/kg | 24 hours | 0.27% |
| Boado et al., 2010 (Boado et al. 2010) | Mouse TfR IgG | Anti-Aβ scFv | ~ 5 nmol/kg | 1 hour | 3.5% total (not capillary depleted) |
| Yu et al., 2011 [30] | Mouse TfR IgG low affinity | BACE1 bispecific antibody in WT mice | ~133 nmol/kg | 12-48 hours | 0.6-0.8% |
| Zuchero et al., 2016 [13] | CD98hc IgG | Anti-BACE1 in WT mice | Tracer (not specified) | 1-48 hours | 1-6% |
| Sehlin et al., 2016 [65] | Mouse TfR IgG 8D3 | mAb158 vs Aβ protofibrils in TgArcSwe mice | ~3 nmol/kg | 4-72 hours | 0.15 - 0.25% |
| Boado & Pardridge 2017 [8] | Insulin R IgG | Iduronidase in rhesus | 0.1 mg/kg | 2 hours | 1.1% ID/100 g brain |
| Hultqvist et al., 2017 [66] | Mouse TfR scFv 8D3 x2 | mAb158 in TgArcSwe mice | 0.25 nmol/kg<br>50 nmol/kg | 72 hours | 2.2%<br>0.33% -0.75% |
| Sonoda et al., 2018 [6] | Human TfR IgG | hIDS in cyno monkeys | 20 nmol/kg | 24 hours | ~0.4% |
| Wicher et al., 2019<br>WO 2019/246288A1 | CD98hc VNAR | Human Fc | 25 nmol/kg | 3-4 hours | 1.3% total (not capillary depleted) |
| Kariolis et al., 2020 [32] | Human TfR binding IgG Fc | BACE 1 IgG in human TfR mice | 300 nmol/kg | 24 hours | ~3.4% |
| Logan et al., 2021 [67] | Human TfR binding IgG Fc | Progranulin in GRN$^{-/-}$ mice | 40 nmol/kg<br>400 nmol/kg | 24 hours | 0.25%<br>0.05% |
| Shin et al., 2022 [12] | Anti-IGF1R scFv | Alpha-synuclein IgG in mice | 340 nmol/kg | 24 hours | 0.4% |
| Van Lengerich et al., 2023 [68] | Human TfR binding IgG Fc | Activating mTREM2 IgG 4D9 in human TfR mice | 66 nmol/kg | 1 day | 1.1-1.5% |
| Gehrlein et al., 2023 [69] | Mouse TfR binding IgG Fab | Recombinant gluco-cerebrosidase in WT and transgenic mice | ~15 nmol/kg | 24 h | ~3% |
| Esparza et al., (this paper) | Mouse TfR nanobody M1<sub>R56H, P96H, Y102H</sub> | 13A7-13A7 nanobodies in WT mice | 30 nmol/kg | 1 hour | 1.7% |
| Esparza et al., (this paper) | Mouse TfR nanobody M1<sub>R56H, P96H, Y102H</sub> | 13A7-13A7-Nb80 nanobodies in WT mice | 1 nmol/kg<br>3 nmol/kg<br>30 nmol/kg | 4 hours | 13.4%<br>8.8%<br>3.5% |

### Supplemental Discussion

The nanobody constructs developed here have relatively fast kinetics and may be potentially useful for short-acting therapeutics and diagnostic applications. However, for applications where a long duration of action is favorable such as treatment of chronic brain diseases, solutions based on receptor mediated transcytosis using conventional immunoglobulins may be preferred [6] [8] [12] [13] [30] [31] [32]. It may be possible to further prolong the duration of action of nanobody-based constructs by making hybrid nanobody-IgG fusions, conjugation to polyethylene glycol, or other approaches, but these may negate many of the advantages of nanobody-based constructs noted elsewhere. Similarly, nanobody-based constructs to not induce IgG Fc-mediated immune responses; this may be an advantage for some applications and a disadvantage for others. Furthermore, for applications where high doses of macromolecules must be delivered to relatively localized targets, focal ultrasound mediated blood brain barrier opening [70] [71] [72] may be preferred over global delivery via brain shuttle systems.

The concentrations of the nanobody constructs in the extracellular space of the brain have not been measured directly. The extracellular concentrations can be estimated based on the mass of the brain tissue and the approximate extracellular fractional volume, under the assumption that the nanobody constructs remain in the extracellular space. As an example:

- Brain concentration in lysates 4 hours after 30 nmol/kg injection of M1_R56H, P96H, Y102H_ -13A7-13A7-Nb80 = 4.5 nM x 1 ml lysate = 4.5 pmol.
- Brain tissue mass ∼200 mg x ∼20% extracellular volume = 40 µl.
- 4.5 pmol/40 µl = 112.5 nM

If a fraction of the nanobody constructs partition into the intracellular space, the concentrations in the extracellular space would be lower. It is also possible that our capillary depletion and lysis procedure results in some loss of brain parenchymal nanobody construct, in which case we would also be underestimating the true brain extracellular concentrations. An approach like cerebral microdialysis with high molecular weight cutoff catheters [73] [74] [75] would be potentially useful for directly measuring the extracellular fluid levels and dynamics of the nanobody constructs in the living brain.

We do not know whether the histidine mutagenesis strategy employed here can be used to improve BBB transcytosis of nanobodies that bind to other receptors such as CD98hc, InsR, or IGF1R. Similarly, its generalizability to affibodies, scFvs, and other binding reagents has not been determined. There are many natural proteins such as transferrin itself with strongly pH dependent binding [5], and several other proteins have been engineered this way [34] [35] [36]. A recent report described highly cooperative pH dependent dissociation of ^64^Cu carrying polymers in the acidic tumor microenvironment, leading to efficient PET imaging of tumor allografts in mice [76].

As noted, the specificity of the 13A7 nanobody has not been fully established. Recent findings indicate that P2X7 receptors are expressed in microglia but not neurons [77]. There appears to be additional P2X7-like immunoreactivity and function in neurons [78] [79] [80], and the most parsimonious explanation for our results may be that 13A7 should be considered reflecting this sort of P2X7-like immunoreactivity. The 13A7 nanobody was primarily selected as a proof-of-concept biological macromolecule cargo with a highly abundant target in wild-type mouse brain for the purposes of assessing our TfR-binding BBB shuttle system. Thus, the specificity of the 13A7 nanobody does not affect the conclusions of the findings presented here.

The fold-increase in BBB crossing imparted by the M1_R56H, P96H, Y102H_ nanobody cannot be directly calculated, but can only be estimated. As noted, we were not able to detect biotinylated M1_AA_-13A7-13A7-Nb80 (which does not bind mTfR) in capillary depleted brain lysates even after injection of 600 nmol/kg. The ELISA lower limit of quantitation was <0.166 nM, indicating <0.007% injected dose/gram of brain tissue. If passive/non-receptor-mediated BBB crossing permits BBB crossing at 0.007% injected dose/gram of brain tissue, this would indicate that 3.5% injected dose/gram of brain tissue represents 500-fold increased BBB crossing. It is difficult to compare our results indicating <0.007% injected dose/gram of brain tissue with previously reported values of 0.1 to 0.01% injected dose/gram of brain tissue for non-specific BBB crossing of biological macromolecules [1] [2]. First, prior studies typically used immunoglobulins or other proteins rather than nanobody constructs. Second, as noted above and shown in **Suppl. Fig. 2A**, capillary depletion based on quantitative Claudin 5 ELISA was ∼99% complete for our studies whereas the quantitatively completeness of capillary depletion has not usually been reported for studies of this type. Third, it is not known whether 4 hours is the optimal time to measure passive/non-receptor-mediated BBB crossing. Nonetheless, based on our results, it is possible that the BBB is even tighter than previously recognized in healthy young adult mice. BBB dysfunction in aged mice and in mouse models of neurological disorders may instead be partially compromised [81]. It is not known whether partial BBB compromise is related to the reported central nervous system effects of peripherally administered macromolecules in preclinical therapeutic studies.

